# A redox-dependent mechanism for AMPK dysregulation interrupts metabolic adaptation of cancer under glucose deprivation

**DOI:** 10.1101/2021.03.10.434352

**Authors:** Younghwan Lee, Yoko Itahana, Choon Chen Ong, Koji Itahana

## Abstract

Accelerated aerobic glycolysis is a distinctive metabolic property of cancer cells that confers dependency on glucose for survival. However, the selective therapeutic targeting of this vulnerability has yielded mixed results owing to the different sensitivities of each cancer type to glucose removal and lack of insight into the underlying mechanisms of glucose deprivation-induced cell death.

Here, we screened multiple cell lines to determine their sensitivities to glucose deprivation and found that the cell lines most sensitive to glucose deprivation failed to activate AMPK, a major regulator of metabolic adaptation, resulting in metabolic catastrophe.

Notably, glucose deprivation-induced AMPK dysregulation and rapid cell death were observed only in cancer cell lines with high expression of cystine/glutamate antiporter xCT. While this phenomenon was prevented by pharmacological or genetic inhibition of xCT, overexpression of xCT sensitized resistant cancer cells to glucose deprivation. We found that cystine uptake and glutamate export through xCT under glucose deprivation contributes to rapid NADPH depletion, leading to the collapse of the redox system, which subsequently inactivates AMPK by inhibitory oxidation and phosphorylation. This AMPK dysregulation caused a failure of metabolic switch to fatty acid oxidation upon glucose deprivation, resulting in mitochondrial dysfunction and cell death.

Taken together, these findings suggest a novel cross-talk between the metabolic and signal transduction and reveal a metabolic vulnerability in xCT-high expressing cancer cells to glucose deprivation and provide a rationale for targeting glucose metabolism in these cancer cells.

## Introduction

Cancer cells exhibit a metabolic reprogramming to support increased demand for energy and building blocks for proliferation^1^. Accelerated aerobic glycolysis is a distinctive metabolic property of cancer cells compared to untransformed cells, which is also known as the Warburg effect. Given that glucose is a major nutrient which is required to maintain biosynthetic, bioenergetics and redox homeostasis, this altered metabolism renders cancer cells highly dependent on glucose for viability. Therefore, this property provides unique vulnerabilities for the selective targeting of such glucose-dependent cancer cells^2^.

Previously, we and others have shown that while some cancer cells demonstrate strict dependency on glucose for their survival, undergoing rapid cell death in a matter of hours following glucose deprivation, others can adapt to glucose starvation, similar to that of untransformed cells^3, 4^. Tissue origin and oncogenic mutations generate a variety of metabolic reprogramming by different means and accordingly confer different dependence on glycolysis. Owing to the lack of known markers to indicate the degree of glucose-dependence, the outcome of glucose restriction strategies for cancer therapy is not always predictable or successful. Therefore, an establishment of biomarkers for the prediction of treatment sensitivity and resistance for glycolysis-targeting cancer therapy would be necessary for a better outcome.

Several studies have shown that glucose deprivation triggers apoptosis and/or necrosis, or autophagy-related cell death^5^. However, the underlying mechanism of glucose deprivation-induced cell death is still unclear, owing to the differences in cell type and experimental conditions in those studies. Although the underlying mechanism of glucose deprivation-induced cell death has been suggested to involve depletion of adenosine triphosphate (ATP) or damage from an accumulation of reactive oxygen species (ROS), recent evidence indicates the involvement of signaling transduction to regulate cell survival and death in response to metabolic stress^6^. Therefore, the understanding of the cross-talk between the metabolic perturbation and signal transduction that leads to cancer cell death in glucose-dependent cancer would be essential for the rational design of combination therapies to improve efficacy or overcome resistance.

5’ AMP-activated protein kinase (AMPK) is a highly conserved serine/threonine kinase that plays a central role in maintaining cellular metabolic balance. AMPK regulates cellular adaptation to energetic stress by engaging appropriate cellular response to cope with metabolic perturbations and its activation requires Thr172 phosphorylation of the activation loop in the kinase domain of the catalytic α subunit by upstream kinases such as liver kinase B1 (LKB1), calcium/calmodulin-dependent protein kinase 2 (CAMKK2)^8, 9^ and TGFβ-activated kinase 1 (TAK1)^10^. Upon activation by starvation, AMPK restores metabolic homeostasis by stimulating ATP-regenerating catabolic processes such as induction of fatty acid oxidation (FAO)^4^, autophagy^11, 12^, and mitochondrial homeostasis^13, 14^ while attenuating anabolic ATP-consuming biosynthetic processes such as suppression of the target of rapamycin complex 1 (TORC1) signaling-mediated protein synthesis^15–17^ and lipid synthesis^4^.

Therefore, proper AMPK activation is required for overcoming metabolic stress. For instance, AMPK protects leukemia-initiating cells from metabolic stress^18^, promotes cancer metastasis^19^ and mediates anoikis resistance^20^, whereas LKB1-null cancers are highly vulnerable to energetic stress due to failure of metabolic adaptation^4, 21^. Thus, understanding different AMPK regulation of glucose addicted versus resistant cancer cells under metabolic stress would be necessary for gaining insight for glucose metabolism targeting therapeutic intervention.

Here, we screened multiple cancer cell lines to determine their sensitivity to glucose deprivation. We identified that cancer cell lines highly sensitive to glucose deprivation express high levels of cystine/glutamate antiporter xCT (xCT), by which expression levels determine the sensitivity to glucose deprivation by affecting intracellular NADPH levels under glucose deprivation. Intracellular redox system collapse causes AMPK dysregulation via inhibitory oxidation and phosphorylation, which renders these glucose-dependent cancer cells unable to adapt to glucose deprivation. Therefore, we suggest xCT can be a biomarker for glucose metabolism targeting cancer therapy and NADPH depletion is the metabolic determinant for glucose deprivation-induced cell death. Also, our data reveal a previously underappreciated crosstalk between metabolism and signaling transduction under metabolic stress, leading to cancer cell death.

## Results

### Cancer cell lines display different AMPK-mTORC1 signaling in response to glucose deprivation

To investigate the signaling mechanisms underlying glucose deprivation-induced cell death, we screened multiple cell lines to determine their sensitivity to glucose deprivation. We incubated cells with glucose-free media for the indicated period (0, 9 or 18 hours) and measured cell death using propidium iodide (PI) exclusion assay. As serum contains a very low amount of glucose, we cultured cells in media with a dialyzed serum which has the necessary growth factors but retains undetectable levels of glucose. Two cancer cell lines (U2OS and U251MG; which will be referred to as “sensitive cell lines”) showed the highest sensitivity to glucose deprivation by showing rapid loss of cellular adhesion within 3 hours, followed by PI-positive cell death within 9 hours. The other two cancer cell lines (SW480 and MCF7) showed a modest response to glucose deprivation with cell death within 18 hours (which will be referred to as “intermediate sensitive cell lines”). The remaining cancer cell lines (A375 and H1299), kidney embryonic epithelial cell line (HEK293T) and normal fibroblast cell lines (WI-38 and IMR-90) remained viable under glucose deprivation (which will be referred to as “resistant cell lines”) (Fig. 1A and Supplementary Fig. S1A). Next, we investigated whether glucose deprivation-induced cell death is a regulated form of cell death. The media for culturing U2OS cells were changed from 25 mM glucose to 0 mM or 1 mM which mimics low glucose concentration in the physiological tumor environment^22, 23^. While U2OS cells cultured in 1 mM glucose maintained viability, cells under glucose deprivation lost viability at 9 hours after glucose withdrawal (Fig. 1B). Interestingly, the cell death in U2OS cells was neither apoptosis, necroptosis nor ferroptosis as none of the inhibitors of regulated cell death [apoptosis inhibitor (Z-VAD-FMK), necroptosis inhibitors (necrostatin-1 and necrosulfonamide) and iron chelator (deferoxamine)] were able to prevent glucose deprivation-induced cell death (Fig. 1B and Supplementary Fig. S1B).

**Figure 1.**
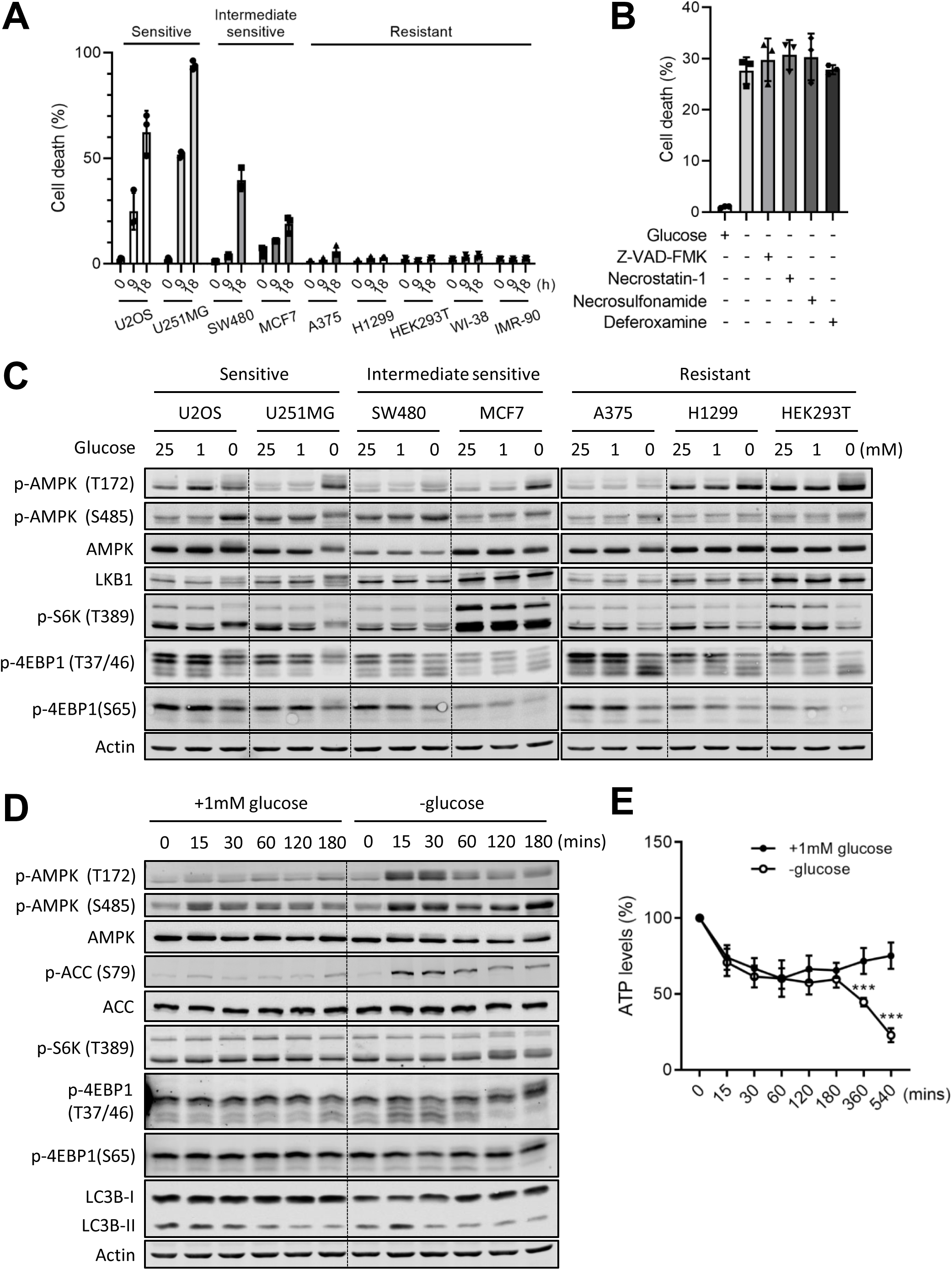
Different sensitivity and AMPK regulation under glucose deprivation in multiple cell lines. (A) Propidium iodide (PI) exclusion assay was performed using the indicated cell lines at the indicated time points after glucose deprivation (0, 9, or 18 hours). (B) PI exclusion assay was performed using U2OS cells cultured in media with or without 1 mM glucose for 9 hours. The indicated cell death inhibitor was treated simultaneously. (C) Western blotting analysis was performed using the indicated cell lines cultured in 25, 1, or 0 mM glucose for 3 hours. (D) Time-course analysis of Western blotting was performed using U2OS cells cultured with or without 1 mM glucose. (E) Time-course analysis of intracellular adenosine 5’-triphosphate (ATP) levels was performed using U2OS cells cultured with or without 1 mM glucose. The mean percentage ± SD of three independent experiments is shown. Images of Western blotting analysis are representative of three independent experiments. Unpaired two-tailed Student’s t-test was performed for statistical analysis, ***P<0.001.

AMPK and mTOR are the central regulators of cellular response to metabolic stress (Supplementary Fig. S1C) ^24^. AMPK senses energy depletion and is activated to maintain energy homeostasis during glucose deprivation^25, 26^. We asked how these central metabolic checkpoints are regulated in sensitive and resistant cell lines. We incubated multiple cell lines in media with different glucose concentrations [25 mM (standard glucose concentration in DMEM), 1 mM (low glucose) or 0 mM (no glucose)] for 3 hours and performed Western blotting analysis (Fig. 1C). In U2OS cells, Thr172 phosphorylation of AMPK that is required for AMPK activation^7^, increased when incubated in 1 mM glucose compared to 25 mM glucose. However, Thr172 phosphorylation under 0 mM glucose was lower than that of 1 mM glucose. AMPK is also phosphorylated in the serine/threonine-rich loop (ST loop) which negatively regulates its kinase activity^27, 28^. Interestingly, one of such inhibitory phosphorylation at the Ser485 of AMPK was increased under 0 mM glucose compared to 1 mM glucose. Besides, U2OS showed an electrophoretic mobility shift of total AMPK, which indicates the possible involvement of hyper-phosphorylation or other post-translational modification of multiple residues. An electrophoretic mobility shift of total AMPK and phosphorylated AMPK was also observed in U251MG cells, another sensitive cell line. In contrast, the intermediate sensitive and resistant cell lines showed no electrophoretic mobility shift of total AMPK, but increased or mildly increased Thr172 phosphorylation of AMPK in glucose-free condition.

To further investigate the activation state of AMPK, we examined mTORC1 signaling, another nutrient sensor that controls cell growth, proliferation and autophagy. When nutrient availability is limited, mTORC1 is inactivated, in part through AMPK activation^15, 16^. Interestingly, while the intermediate sensitive and resistant cell lines showed decreased mTORC1 activity under glucose deprivation (0 mM glucose), as indicated by the changes in phosphorylation pattern of S6K (Thr389) and 4EBP1 (Thr37/46 and Ser65) as described elsewhere^29^, the sensitive cell lines (U2OS and U251MG) maintained high mTORC1 activity, as indicated by the increased molecular weight of phosphorylation of S6K and 4EBP1 as described^29^. These data indicate that sensitive cell lines maintained high mTOR and low AMPK activities under glucose deprivation, whereas resistant cell lines increased AMPK and reduced mTOR activities to adapt to the metabolically challenging conditions.

One thing to highlight is that all the cell lines used in this study expressed LKB1 which is an upstream kinase of AMPK Thr172. Only the sensitive cell lines showed a clear electrophoretic shift of LKB1, which may represent a highly phosphorylated status of LKB1 under glucose deprivation.

To gain a better insight into how the AMPK-mTORC1 signaling is dysregulated in sensitive cell lines upon glucose deprivation, we examined the above phosphorylations in a time-dependent manner in U2OS cells (Fig. 1D). Under glucose deprivation, Thr172 phosphorylation of AMPK was increased almost immediately after glucose withdrawal, but was diminished around the 1-2 hour time point. Similarly, a well-characterized AMPK substrate, acetyl-CoA carboxylase (ACC) was also phosphorylated at Ser79 immediately upon glucose deprivation and was diminished subsequently while total ACC amounts were unchanged. The Ser485 inhibitory phosphorylation of AMPK was continuously increased under glucose deprivation. Furthermore, high mTORC1 activity was maintained, as indicated by phosphorylation patterns of 4EBP1 and S6K. The LC3B-II/LC3B-I ratio, the hallmark of autophagy which is positively regulated by AMPK and negatively regulated by mTORC1^11, 12^, was slightly decreased. These results suggest that after experiencing the transient AMPK activation by glucose withdrawal, U2OS cells maintained low AMPK and high mTORC1 activities. A similar trend was observed in another sensitive cell line, U251MG cells (Supplementary Fig. S1D). These data indicate that the sensitive cell lines display dysregulated AMPK-mTORC1 signaling under glucose deprivation and maintain the phosphorylation signature of a ‘fed’ state while under starvation conditions.

To investigate whether the AMPK-mTORC1 signaling dysregulation was due to altered energy status, we measured intracellular ATP levels in U2OS cells in a time-dependent manner (Fig. 1E). U2OS cells cultured in both 1 and 0 mM glucose showed similarly decreased ATP levels up to 3 hours when the initial stage of cell death (cell detachment) occurred in 0 mM glucose, but not in 1mM glucose (Supplementary Fig. S1B). While cells in 1 mM glucose maintained ATP levels, cells in the 0 mM glucose showed further depletion of ATP levels after 3 hours. These data indicate that energy status does not reflect the dysregulated AMPK-mTORC1 signaling under glucose deprivation and that decreased levels of ATP are not the main trigger of cell detachment from the culture plate.

### Dysregulated AMPK pathway contributes to glucose deprivation-induced cell death in sensitive cell line

Given that AMPK activation is essential to overcome metabolic stress^4, 21^, we hypothesized that dysregulated AMPK signaling in sensitive cell lines underlies the failure to adapt to glucose deprivation. Treatment with either A769662, an AMPK allosteric activator^30^ or 2-deoxyglucose (2-DG), a non-metabolizable glucose analog and indirect AMPK activator^31^, rescued glucose withdrawal-induced cell death in U2OS cells (Fig. 2A and B and Supplementary Fig. S2A and B), suggesting that AMPK activation is sufficient to rescue cell death induced by glucose deprivation. High AMPK activity by both A769662 and 2-DG treatment under glucose deprivation was confirmed by Western blotting analysis, showing increased Thr172 phosphorylation of AMPK and phosphorylation of ACC (Fig. 2C and Supplementary Fig. S2C). AMPK activation by these treatments decreased mTORC1 activity, as indicated by phosphorylation patterns of 4EBP1 and S6K (Fig. 2C and Supplementary Fig. S2C). A similar phenotype was observed in U251MG cells (Supplementary Fig. S2F). These data suggest that glucose deprivation-induced cell death in sensitive cell lines is due to blunted AMPK activation.

**Figure 2.**
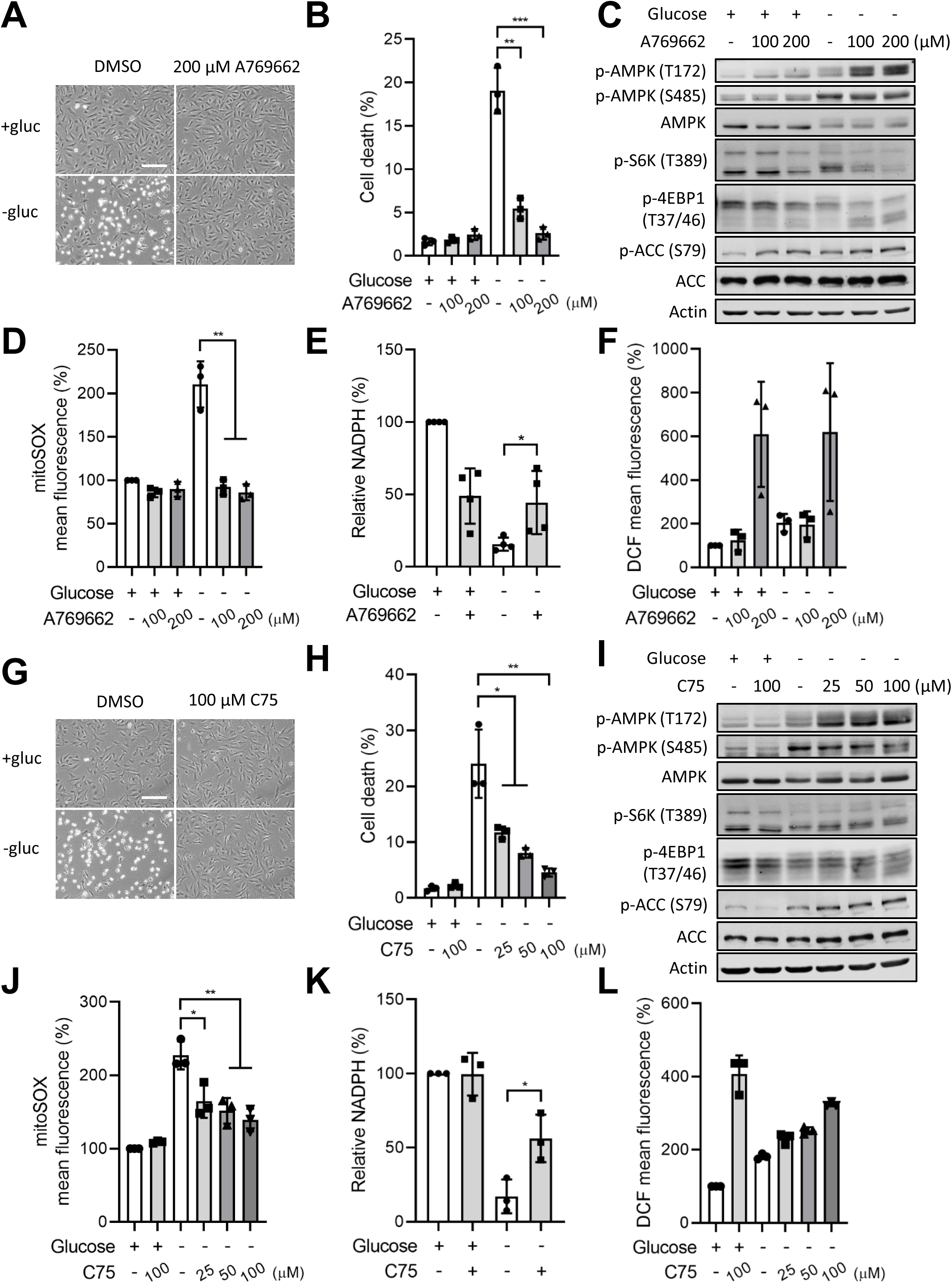
AMPK dysregulation contributes to glucose deprivation-induced cell death. U2OS cells in media with or without 1 mM glucose were treated with indicated doses of A769662 for 3 hours and representative images were taken using phase-contrast microscopy (A); Propidium iodide (PI) exclusion assay was performed at 9 hours (B); Western blotting analysis was performed at 3 hours (C); mitochondrial ROS was measured at 3 hours (D); intracellular NADPH levels were measured at 2 hours (E); cytosolic ROS was measured at 3 hours (F). U2OS cells in media with or without 1 mM glucose treated with indicated doses of C75 for 3 hours and representative images were taken using phase-contrast microscopy (G); PI exclusion assay was performed after 9 hours (H); Western blotting analysis was performed at 3 hours (I); mitochondrial ROS was measured at 3 hours (J); intracellular NADPH levels were measured at 2 hours (K); cytosolic ROS was measured at 3 hours (L). Scale bars, 500 μm. The mean percentage ± SD of three or more than three independent experiments is shown. Images of Western blotting analysis are representative of three independent experiments. Unpaired two-tailed Student’s t-test was performed for statistical analysis, *P<0.05, **P<0.01, ***P<0.001.

AMPK maintains catabolic and anabolic metabolic homeostasis by regulating protein synthesis, energy generation, autophagy and redox status^24^. We wondered which functional pathway downstream of AMPK is critical for preventing cell death caused by glucose deprivation. First, we tested whether AMPK protected cancer cells from metabolic stress through mTORC1 inhibition. We treated U2OS cells with rapamycin to inhibit mTORC1 signaling because it rescued metabolic stress-induced cell death in previous studies^17, 20^. If glucose deprivation-induced cell death is due to the defect in mTORC1 inhibition because of blunted AMPK, rapamycin treatment is expected to rescue cell death of U2OS cells in the absence of glucose. However, rapamycin treatment did not affect glucose deprivation-induced cell death (Supplementary Fig. S3A and B), although it completely inhibited mTORC1 signaling as shown by decreased phosphorylation of S6K and decreased molecular weight of phosphorylated 4EBP1 as reported elsewhere^32^ (Supplementary Fig. S3C). These data suggest that glucose deprivation-induced cell death is not because of the lack of mTORC1 inhibition due to the blunted AMPK. We also observed that rapamycin treatment did not increase LC3-II/LC3-I ratio in the absence of glucose as efficiently as in the presence of glucose, indicating that autophagy is prevented under glucose deprivation. This observation is consistent with the idea that AMPK is blunted under glucose deprivation since AMPK promotes autophagy through not only mTORC1 inhibition but also direct phosphorylation of Ulk1^11, 12^.

Glucose is the main source of NADPH which is generated through the pentose phosphate pathway (PPP) and glucose deprivation induces oxidative stress by an accumulation of cytosolic and mitochondrial ROS^4, 6, 33, 34^. Therefore, glucose deprivation-mediated ROS production was considered as a metabolic determinant of glucose deprivation-induced cell death in the previous studies^4, 6, 33, 34^. AMPK maintains redox balance by upregulating NADPH levels and reduced glutathione (GSH) through the inhibition of fatty acid synthesis and promotion of FAO, processes that consume or generate NADPH, respectively via phosphorylating ACC^4, 35^. Therefore, we hypothesized that AMPK activation rescues glucose withdrawal-induced cell death by restoring redox balance via switching from glycolysis to FAO. A769662 treatment prevented mitochondrial ROS accumulation and partially rescued NADPH depletion induced by glucose deprivation (Fig. 2D and E). However, it further increased glucose deprivation-induced cytosolic ROS accumulation (Fig. 2F), which is contrary to the previously suggested causality between cytosolic ROS and cell death under glucose deprivation^4, 6^. On the other hand, the 2-DG treatment prevented glucose deprivation-induced mitochondrial and cytosolic ROS accumulation (Supplementary Fig. S2D and E). Together, these data suggest that the mitochondrial redox status and intracellular NADPH depletion, but not cytosolic ROS, might be directly associated with glucose deprivation-induced cell death.

Next, to determine whether regulation of fatty acid metabolism is sufficient to protect cancer cells from glucose deprivation, we treated U2OS cells with C75, a de novo fatty acid synthesis inhibitor that also facilitates FAO. C75 treatment prevented glucose deprivation-induced cell death (Fig. 2G and H), suggesting that low FAO and high fatty acid synthesis contribute to cell death induced by glucose withdrawal. Interestingly, C75 treatment activated AMPK signaling as indicated by increased phosphorylation of AMPK at Thr172 and ACC, although its effect on mTORC1 signaling was marginal as shown by phosphorylation patterns of S6K and 4EBP1 (Fig. 2I). Also, C75 treatment prevented mitochondrial ROS accumulation (Fig. 2J) and partially restored NADPH depletion (Fig. 2K), although it further increased cytosolic ROS (Fig. 2L), mimicking the effect of the AMPK activator. These results suggest that replenishing NADPH by promoting FAO prevented the collapse of the redox system, thereby preventing dysregulation of AMPK signaling under glucose deprivation.

Together, AMPK activation rescues glucose deprivation-induced cell death mainly by regulating redox homeostasis via FAO. Sensitive cancer cells are not able to switch from glycolysis to FAO due to AMPK dysregulation, preventing adaptation to metabolic stress.

### Mitochondrial dysfunction plays a role in glucose deprivation-induced cell death

Because glucose deprivation-induced mitochondrial ROS accumulation was highly correlated to cell death, we investigated whether mitochondrial dysfunction was the possible cause of cell death. Direct replenishing of the mitochondrial tricarboxylic acid (TCA) cycle with cell-permeable methyl-pyruvate (me-pyruvate) or dimethyl α-ketoglutarate (DMKG) in U2OS cells partially rescued glucose deprivation-induced cell death (Fig. 3A and B), suggesting restoring mitochondrial function contributes to rescuing glucose deprivation-induced cell death. Me-pyruvate or DMKG treatment activated AMPK signaling as indicated by increased phosphorylation of AMPK at Thr172 and ACC although mTORC1 signaling was not affected (Fig. 3C). Also, they prevented mitochondrial ROS accumulation (Fig. 3D) and partially rescued NADPH depletion (Fig. 3E), although neither TCA cycle substrates rescued cytosolic ROS accumulation (Fig. 3F). These data suggest that mitochondrial dysfunction plays a role in glucose deprivation-induced AMPK dysregulation and cell death and confirm no causality between cytosolic ROS and cell death under glucose withdrawal.

**Figure 3.**
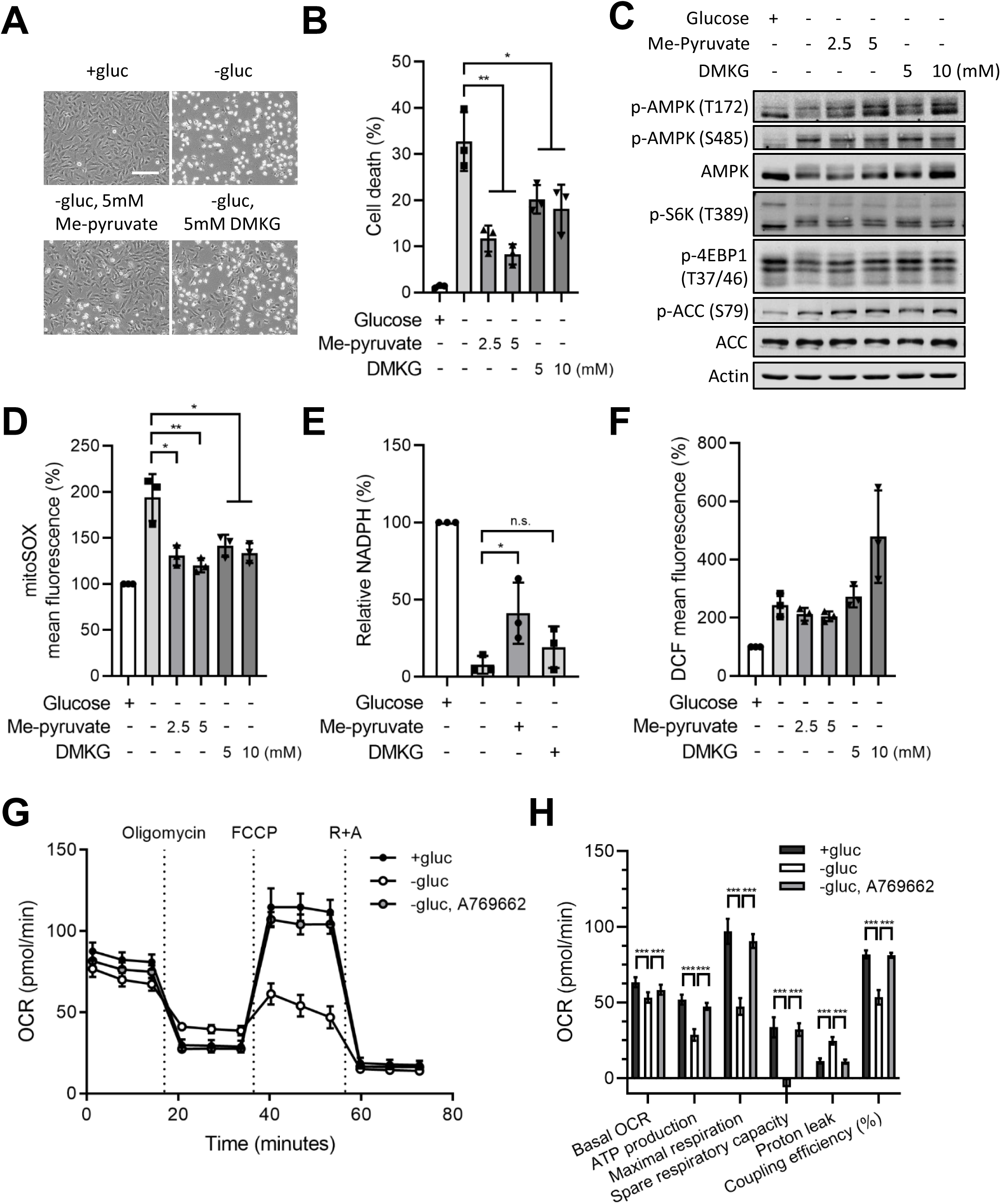
Mitochondrial dysfunction contributes to glucose deprivation-induced cell death. U2OS cells in media with or without 1 mM glucose were treated with indicated doses of methyl-pyruvate (Me-pyruvate) or dimethyl α-ketoglutarate (DMKG) for 3 hours and representative images were taken using phase-contrast microscopy (A); Propidium iodide (PI) exclusion assay was performed at 9 hours (B); Western blotting analysis was performed at 3 hours (C); mitochondrial ROS was measured at 3 hours (D); intracellular NADPH levels were measured at 2 hours (E); cytosolic ROS was measured at 3 hours (F). (G and H) Mitochondrial oxygen consumption rate (OCR) was measured using a Seahorse extracellular flux analyser. U2OS cells were cultured with or without 1 mM glucose and 200 μM A769662 was treated simultaneously. Scale bars, 500 μm. The mean percentage ± SD of three independent experiments is shown. Images of Western blotting analysis are representative of three independent experiments. Unpaired two-tailed Student’s t-test was performed for statistical analysis, *P<0.05, **P<0.01, ***P<0.001, n.s. not significant.

To further investigate the effect of glucose deprivation on mitochondrial dysfunction, we assessed the mitochondrial oxygen consumption rate (OCR) and mitochondrial metabolic parameters by Seahorse assay. Compared to U2OS cells cultured in 1 mM glucose, cells under glucose deprivation showed a decrease in basal mitochondrial OCR, mitochondrial ATP production, maximal respiration, spare respiratory capacity (SRC), and coupling efficiency and an increase in proton leak, all of which could be a sign of mitochondrial damage under glucose deprivation (Fig. 3G and H). A769662 treatment under glucose deprivation rescued these mitochondrial metabolic parameters, implying that replenishing the TCA cycle by accelerating FAO improves mitochondrial fitness. Given that SRC represents mitochondrial fitness as it correlates with bioenergetics adaptability in responding to metabolic stress^36^, glucose deprivation-induced SRC depletion indicates that mitochondria become dysfunctional under glucose withdrawal and that the mitochondrial ROS accumulation could be due to defective mitochondrial function, rather than promotion of mitochondrial oxidative phosphorylation (OXPHOS).

Taken together, these data suggest that: a) glucose deprivation induces mitochondrial dysfunction; b) mitochondrial dysfunction might be due to a lack of TCA cycle substrate; c) AMPK dysregulation plays a role in mitochondrial dysfunction because it could not activate FAO to replenish the TCA cycle.

### Glucose deprivation dysregulates AMPK signaling in a redox-dependent manner

Our next question was how glucose deprivation dysregulates AMPK function in sensitive cell lines. Given that redox status correlates well with cell survival under glucose withdrawal (Fig. 2), we hypothesized that glucose removal-induced oxidative stress dysregulates AMPK signaling, leading to cell death. Treatment with anti-oxidants, N-acetyl-cysteine (NAC) or GSH prevented glucose deprivation-induced cell death (Fig. 4A and B), mitochondrial and cytosolic ROS accumulation (Fig. 4C and D) and NADPH depletion (Fig. 4E) in U2OS cells. NAC or GSH treatment enabled U2OS cells to activate AMPK signaling under glucose deprivation as indicated by increased Thr172 phosphorylation of AMPK and phosphorylation of ACC and decreased Ser485 phosphorylation of AMPK. Also, these treatments decreased mTORC1 activity, as indicated by changes in phosphorylation of 4EBP1 and S6K (Fig. 4F). A similar trend was observed in U251MG cells (Supplementary Fig. S4A). These data indicate that oxidative stress induced by glucose withdrawal perturbs AMPK signaling, leading to a failure in metabolic adaptation.

**Figure 4.**
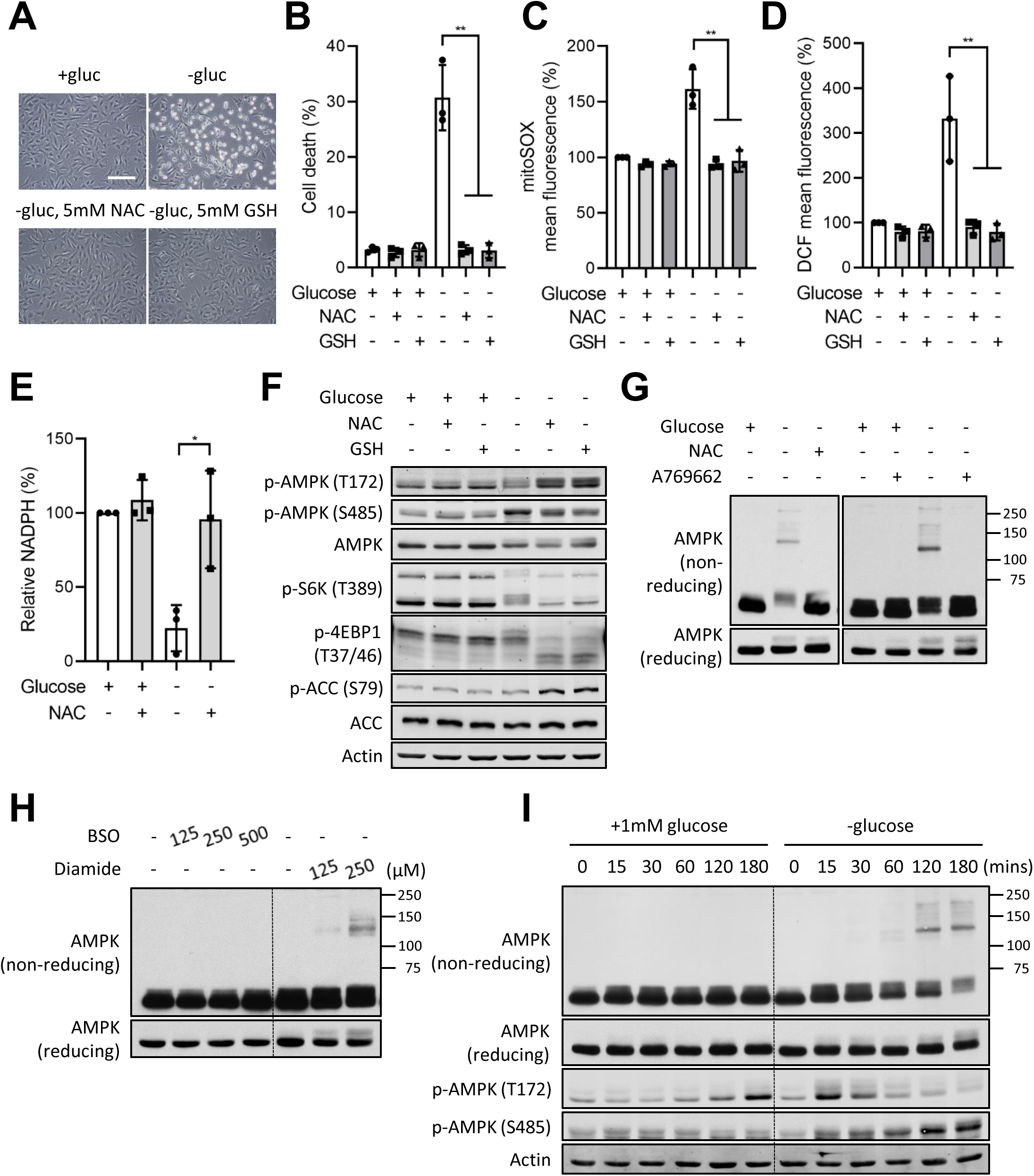
Glucose deprivation dysregulates AMPK signaling in a redox-dependent manner. U2OS cells in media with or without 1 mM glucose were treated with 5 mM N-acetyl-cysteine (NAC) or reduced glutathione (GSH) for 3 hours and representative images were taken using phase-contrast microscopy (A); Propidium iodide (PI) exclusion assay was performed at 9 hours (B); mitochondrial (C) or cytosolic (D) ROS was measured at 3 hours; intracellular NADPH levels were measured at 2 hours (E); Western blotting analysis was performed at 3 hours (F). (G) U2OS cells in media with or without 1 mM glucose were treated with 5 mM NAC or 200 μM A769662 for 3 hours. Non-reducing or reducing Western blotting analysis was performed. (H) U2OS cells in media with 1 mM glucose were treated with indicated doses of buthionine sulfoximine (BSO) for 6 hours or diamide for 2 hours. Non-reducing or reducing Western blotting analysis was performed. (I) Time-course analysis of non-reducing or reducing Western blotting analysis was performed using U2OS cells cultured with or without 1 mM glucose. Scale bars, 500 μm. The mean percentage ± SD of three independent experiments is shown. Images of Western blotting analysis are representative of three independent experiments. Unpaired two-tailed Student’s t-test was performed for statistical analysis, *P<0.05, **P<0.01.

Although ROS-induced oxidative stress was thought to cause nonspecific damage to cellular components such as lipids, DNA and proteins, recent evidence suggests that ROS can modulate protein function by oxidizing specific cysteine residues on target proteins^37, 38^. In particular, intracellular NADPH levels affect protein oxidation because NADPH is essential for recycling two major thiol-dependent antioxidant pathways, GSH/GSH reductase (GSR) and thioredoxin (TXN)/ thioredoxin reductase (TXNRD) which maintain cellular thiol redox homeostasis and protein dithiol/disulfide balance^39^. Both GSH and TXN reduce protein disulfide bonds to dithiols which is essential for maintaining protein function^39^. Therefore, we speculated two possible underlying mechanisms of glucose deprivation-induced blunted AMPK activation: first, AMPK undergoes oxidative modification under glucose deprivation, which inhibits AMPK activity and function^40^. Second, glucose deprivation affects AMPK function through inhibitory phosphorylation by redox-sensitive kinase. To examine whether AMPK is directly oxidized under glucose deprivation, we measured the electrophoretic mobility shift of AMPK in the non-reducing SDS-PAGE which does not contain the reducing agents such as DTT and β-mercaptoethanol (β-ME). Glucose deprivation in U2OS cells induced a shift of molecular weight of AMPK only in the non-reducing SDS-PAGE, which was prevented by NAC or A769662 treatment. Different levels of mobility shift of AMPK were observed under glucose deprivation. It may be due to different extent of oxidation of protein. The mobility shift of AMPK was not observed in reducing SDS-PAGE (Fig. 4G). Similar results were obtained in U251MG cells (Supplementary Fig. S4B). These results suggest that glucose deprivation induces aggregation of AMPK through protein disulfide bond formation. To determine whether depletion of NADPH is required for AMPK oxidation, we administered different pro-oxidants, diamide or buthionine sulfoximine (BSO) to U2OS cells and examined AMPK oxidation. Diamide oxidizes thiol-containing proteins, leading to depletion of NADPH. BSO is an inhibitor of γ-glutamylcysteine synthetase which induces ROS by inhibiting the synthesis of GSH, but does not deplete NADPH immediately^41^. Diamide treatment was sufficient to induce AMPK oxidation in non-reducing SDS-PAGE, whereas BSO treatment did not induce AMPK oxidation although both pro-oxidant reagents cause accumulation of cytosolic ROS (Fig. 4H). These data indicate that intracellular NADPH level determines the AMPK oxidation status and sensitivity to glucose deprivation.

Next, to understand the sequential order of glucose deprivation-induced oxidation and aberrant phosphorylation of AMPK, we measured oxidation and phosphorylation of AMPK in a time-dependent manner (Fig. 4I). We observed an inverse correlation between Thr172 phosphorylation and inhibitory Ser485 phosphorylation with increasing oxidation of AMPK in a time-dependent manner, indicating that inhibitory oxidation and inhibitory phosphorylation of AMPK could negatively co-regulate the function of AMPK together.

Taken together, these data suggest that glucose deprivation-induced cell death is caused by the depletion of NADPH and that NADPH could be a major metabolic determinant of the oxidation status of redox-sensitive proteins such as AMPK under glucose deprivation.

### PKC and GSK3 induces inhibitory phosphorylation of AMPK under glucose deprivation

Since glucose deprivation induced an electrophoretic mobility shift of total AMPK in sensitive cell lines (Fig. 1C), we tested whether this was due to aberrant hyper-phosphorylation of AMPK. Incubation of cell lysate with λ phosphatase, a serine/threonine/tyrosine phosphatase reversed the glucose deprivation-induced AMPK mobility shift, indicating that the mobility shift was due to phosphorylation (Fig 5A).

**Figure 5.**
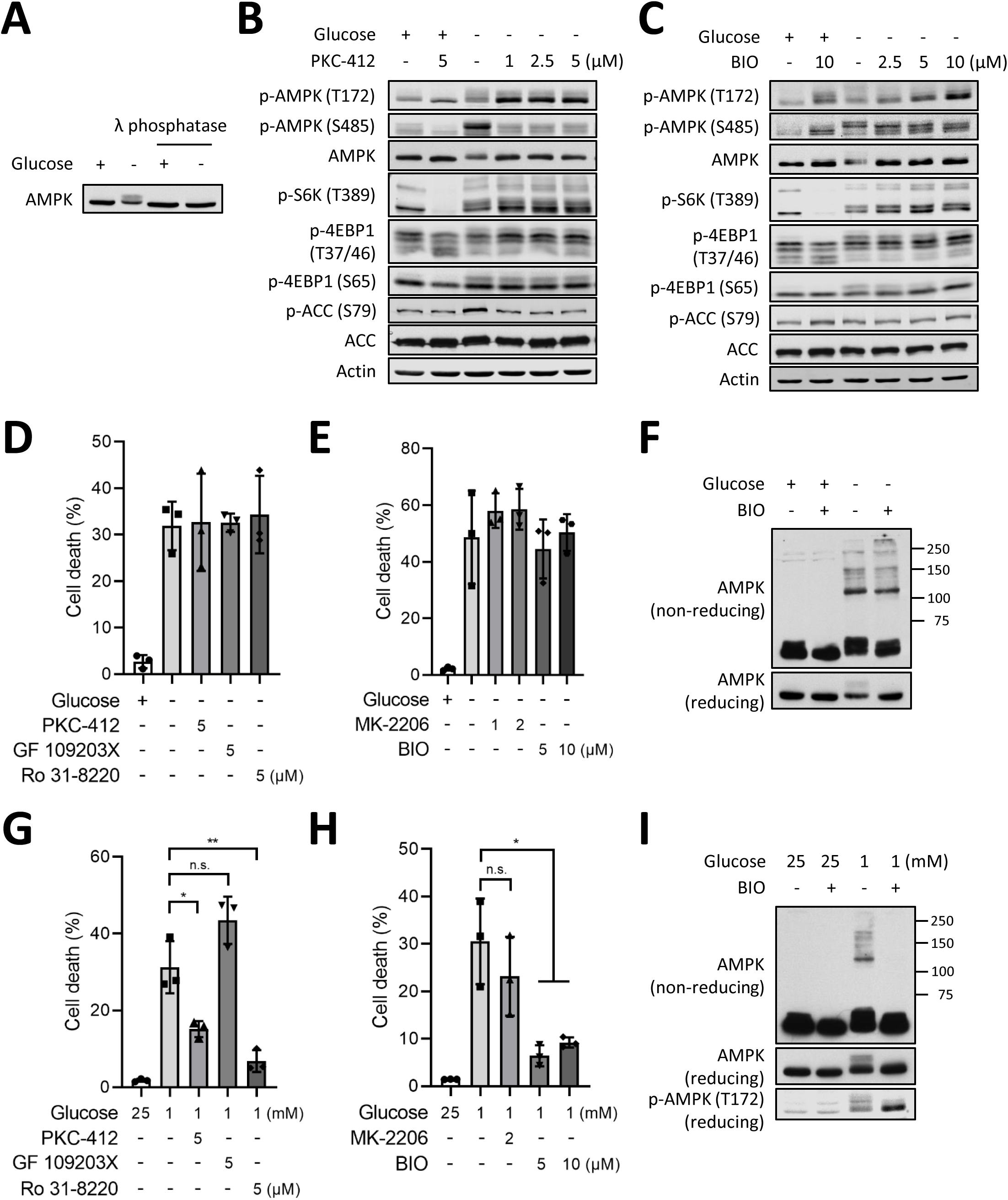
The role of inhibitory phosphorylation of AMPK under glucose deprivation. (A) U2OS cells were cultured with or without 1 mM glucose for 3 hours and harvested. λ phosphatase-digested lysates were used for Western blotting analysis. (B and C) U2OS cells were treated with indicated doses of PKC-412 or BIO in media with or without 1 mM glucose for 3 hours and harvested. Western blotting analysis was performed. (D and E) Propidium iodide (PI) exclusion assay was performed using U2OS cells cultured in media with or without 1 mM glucose for 9 hours. The indicated kinase inhibitor was treated simultaneously. (F) U2OS cells were treated with 5 μM BIO for 3 hours in the media with or without 1 mM glucose and harvested. Non-reducing or reducing Western blotting analysis was performed. (G and H) PI exclusion assay was performed using U2OS cells cultured in media with 25 or 1 mM glucose for 24 hours. The indicated kinase inhibitor was treated simultaneously. (I) U2OS cells were treated with 5 μM BIO for 16 hours in the media with 25 or 1 mM glucose and harvested. Non-reducing or reducing Western blotting analysis was performed. The mean percentage ± SD of three independent experiments is shown. Images of Western blotting analysis are representative of three independent experiments. Unpaired two-tailed Student’s t-test was performed for statistical analysis, *P<0.05, **P<0.01, n.s. not significant.

AMPK has multiple phosphorylation residues in the ST loop, such as Ser485 and Thr479, which negatively regulates the function of AMPK. Akt^27, 42^, PKC^43^, and S6K^44^ are known to phosphorylate Ser485, whereas GSK3 is known to phosphorylate Thr479 and sequential phosphorylation sites^28, 45^. Therefore, we hypothesized that glucose deprivation activates a redox-sensitive kinase which inhibits AMPK by phosphorylating Ser485 or Thr479 residues in the ST loop, leading to failure of metabolic adaptation. We chose several kinase inhibitors which are known to regulate inhibitory phosphorylation of AMPK and tested which kinase is responsible for glucose deprivation-induced inhibitory phosphorylation of AMPK. Akt was previously reported as being responsible for Ser485 phosphorylation of AMPK under multiple conditions^27, 42^. However, MK-2206, an Akt inhibitor did not inhibit glucose deprivation-induced Ser485 phosphorylation and did not recover Thr172 phosphorylation and mobility shift of AMPK in U2OS cells (Supplementary Fig. 5A), indicating that Akt is not the main kinase of Ser485 phosphorylation following glucose withdrawal. The treatment with PKC inhibitors, PKC-412, GF 109203X and Ro 31-8220 prevented glucose deprivation-induced inhibitory Ser485 phosphorylation and total AMPK mobility shift and restored Thr172 phosphorylation of AMPK under glucose deprivation in U2OS cells (Fig. 5B and Supplementary Fig. S5B and C), suggesting that PKC might be responsible for glucose deprivation-induced Ser485 phosphorylation. On the other hand, a GSK3 inhibitor, BIO treatment only reversed the glucose deprivation-induced Thr172 phosphorylation of AMPK and total AMPK mobility shift but maintained Ser485 phosphorylation (Fig. 5C). This is in line with a previous study suggesting that GSK3 phosphorylates Thr479 and neighboring residues, but not Ser485^28^. Ser485 phosphorylation is required for and is followed by GSK3-mediated Thr479 phosphorylation and that of neighboring residues^28^. Therefore, PKC-induced Ser485 phosphorylation seems to precede GSK3-mediated phosphorylation. Although PKC and GSK3 inhibitors rescued the AMPK phosphorylation status under glucose deprivation, they did not affect the phosphorylation statuses of 4EBP1, S6K, and ACC, which are downstream of AMPK (Fig. 5B and C and Supplementary Fig. S5B and C). Besides, neither treatment in U2OS cells rescued glucose deprivation-induced cell death (Fig. 5D and E). These data indicate that AMPK might be no longer coupled with its substrates under glucose deprivation and suggest that changing the phosphorylation of AMPK to an active state is not sufficient to makes AMPK fully functional.

Given that protein oxidation can affect its subcellular localization and interactions^38^, we hypothesized that the effects of PKC or GSK3 inhibitors were not enough to overcome glucose deprivation-induced oxidative stress because NADPH levels are depleted too rapidly within 2 hours after glucose withdrawal (Fig. 2E). In the non-reducing SDS-PAGE, we found that GSK3 inhibition did not completely prevent glucose deprivation-induced AMPK oxidation although it prevented AMPK mobility shift induced by phosphorylation in the reducing SDS-PAGE (Fig. 5F), indicating that oxidation modification could be the underlying mechanism that uncouples AMPK from its substrates. To confirm this idea, we employed the low glucose condition (1 mM glucose) where glucose starvation-induced NADPH depletion and cell death occur slowly.

Interestingly, two PKC inhibitors (PKC-412 and Ro 31-8220) and GSK3 inhibitor (BIO) significantly rescued cell death that was measured at 24 hours after low glucose incubation (Fig. 5G and H). Why GF 109203X treatment did not rescue cell death is unclear. But, it might be due to the different selectivity or sensitivity on different PKC isoforms, compared to the other two PKC inhibitors. Further investigation is needed to figure out which PKC isoform is responsible for Ser485 phosphorylation of AMPK under glucose deprivation. These data suggest that glucose deprivation-induced aberrant phosphorylation in the ST loop negatively regulates AMPK function, leading to failure of metabolic adaptation. Consistent with this data, BIO treatment completely prevented low glucose-induced AMPK oxidation in the non-reducing SDS-PAGE and recovered total AMPK mobility shift and Thr172 phosphorylation of AMPK in the reducing SDS-PAGE (Fig. 5I). Taken together, our data reveal that glucose deprivation induces aberrant phosphorylation in the ST loop of AMPK by PKC and GSK3, which inhibits AMPK-mediated adaptive response to metabolic stress.

### Cystine flux is required for glucose deprivation-induced redox collapse, AMPK dysregulation and cell death

Next, we asked whether glucose deprivation alone is sufficient to induce cell death or other nutrients such as amino acids are involved. We cultured U2OS cells in DMEM or amino acid-free DMEM (DMEM-AA) media with and without 1 mM glucose and evaluated PI-positive cell death. As a result, glucose deprivation in DMEM-AA did not induce cell death. However, when essential amino acids (EAA), but not non-essential amino acids (NEAA), were supplemented in the DMEM-AA media, glucose deprivation was able to induce cell death. To further examine which essential amino acid was required for glucose deprivation-induced cell death, we supplemented DMEM-AA with EAAs separately and measured cell death and found that cystine was required (Fig. 6A).

**Figure 6.**
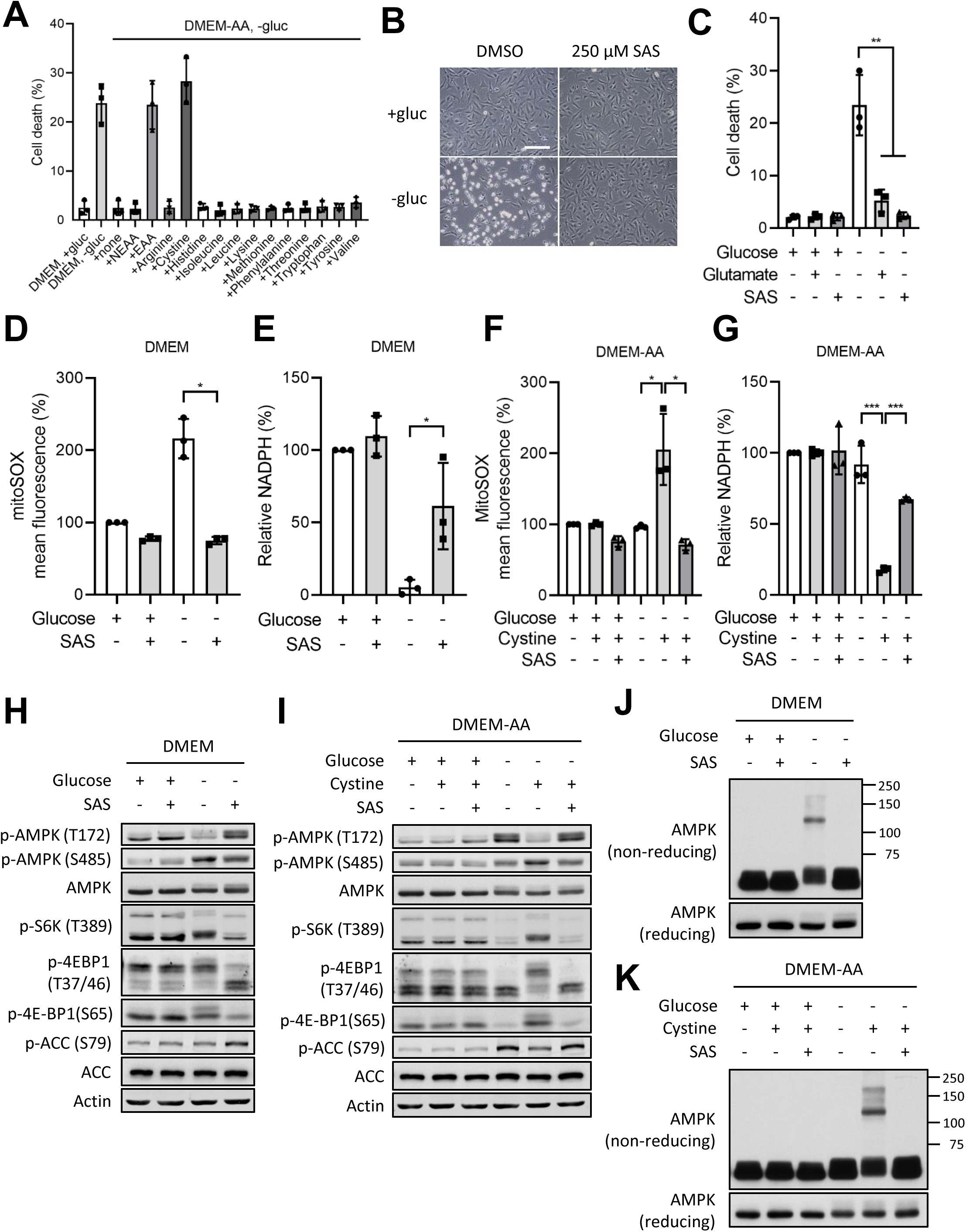
Cystine/glutamate antiporter xCT-mediated cystine uptake is required for glucose deprivation-induced AMPK dysregulation and cell death. (A) Propidium iodide (PI) exclusion assay was performed using U2OS cells cultured in DMEM or amino acid free-DMEM (DMEM-AA) with or without 1 mM glucose for 9 hours. The indicated amino acid was supplemented simultaneously. U2OS cells in the indicated media with or without 1 mM glucose were treated with 250 μM sulfasalazine (SAS) or 3 mM glutamate for 3 hours and representative images were taken using phase-contrast microscopy (B); PI exclusion assay was performed at 9 hours (C); mitochondrial ROS was measured at 3 hours (D and F); intracellular NADPH levels were measured at 2 hours (E and G); Western blotting analysis was performed at 3 hours (H and I); non-reducing or reducing Western blotting analysis was performed at 3 hours (J and K). Scale bars, 500 μm. The mean percentage ± SD of three independent experiments is shown. Images of Western blotting analysis are representative of three independent experiments. Unpaired two-tailed Student’s t-test was performed for statistical analysis, *P<0.05, **P<0.01, ***P<0.001.

Cystine is uptaken through xCT also known as SLC7A11, the light chain of the cystine/glutamate antiporter system x_c_^-^. xCT exchanges intracellular glutamate for extracellular cystine which is rapidly converted to cysteine using NADPH and serves as the precursor for GSH generation^46^. Because GSH is a powerful antioxidant, cancer cells often upregulate xCT to maintain high GSH levels, which contributes to chemoresistance, tumor invasion and poor survival^47–49^. To determine whether cystine uptake through the xCT is required for glucose deprivation-induced cell death, we treated U2OS cells with sulfasalazine (SAS) or extracellular glutamate to inhibit xCT activity. Both xCT inhibitors prevented glucose deprivation-induced cell death (Fig. 6B and C), indicating that the activity of xCT is required for glucose withdrawal-induced cell death. We proceeded to examine whether cystine uptake through xCT is responsible for glucose deprivation-induced redox system collapse. xCT inhibition with SAS in U2OS cells prevented glucose withdrawal-induced mitochondrial ROS accumulation (Fig. 6D) and partially rescued NADPH depletion (Fig. 6E). However, SAS treatment did not prevent cytosolic ROS accumulation (Supplementary Fig. 6A and B), consistent with our earlier data indicating cytosolic ROS was not the determinant of glucose deprivation-induced cell death. In DMEM-AA, glucose deprivation alone was not sufficient to induce mitochondrial ROS accumulation and NADPH depletion. Cystine supplementation in the media was required for these phenotypes which were prevented by SAS treatment (Fig. 6F and G). Given that NADPH is used for recycling oxidized glutathione (GSSG) to the reduced glutathione (GSH), we examined whether GSH metabolism is also regulated by glucose availability and xCT activity. Glucose deprivation in DMEM decreased the GSH/GSSG ratio which was partially restored by xCT inhibition (Supplementary Fig. S6C). In DMEM-AA, glucose deprivation alone was not sufficient to oxidize GSH and cystine supplementation was required to induce complete oxidation of GSH and was partially restored by xCT inhibition (Supplementary Fig. S6D).

Next, we investigate whether xCT-mediated cystine uptake under glucose deprivation dysregulates AMPK signaling. In DMEM, xCT inhibition with SAS in both U2OS maintained high AMPK and low mTORC1 activities under glucose deprivation, as represented by increased phosphorylation of ACC and decreased phosphorylation of 4EBP1 and S6K (Fig. 6H). Importantly, glucose deprivation alone in DMEM-AA activated AMPK signaling, as indicated by increased Thr172 phosphorylation of AMPK and phosphorylation of ACC without change of Ser485 inhibitory phosphorylation of AMPK. Also, it inhibited mTORC1 signaling, as indicated by decreased phosphorylation of S6K and 4EBP1, suggesting that AMPK signaling responding metabolic stress is intact in U2OS cells (Fig. 6I). However, when cystine was supplemented in DMEM-AA under glucose deprivation, AMPK lost its activity. Similar results were obtained in U251MG cells (Supplementary Fig. S6E and F). In the non-reducing SDS-PAGE, AMPK oxidation induced by glucose deprivation in DMEM was prevented by xCT inhibition (Fig. 6J) and glucose deprivation alone in DMEM-AA did not oxidize AMPK. When cystine was supplemented in DMEM-AA, AMPK oxidation was observed under glucose deprivation in U2OS cells (Fig. 6K).

Collectively, our data suggest that glucose deprivation-induced redox system collapse and AMPK dysregulation are due to cystine uptake through xCT. Conversion of cystine to cysteine may accelerate NADPH consumption only under glucose deprivation where NADPH supply is limited. In contrast, when NADPH is in surplus by PPP in the presence of glucose, high cystine uptake through xCT no longer depletes intracellular NADPH and ensues that the enhanced GSH generation can be beneficial for cancer cell survival.

### Sensitive cell lines expressing high levels of xCT show rapid NADPH depletion upon glucose deprivation

Given that the activity of xCT is required for high sensitivity to glucose deprivation, we sought to investigate whether there is a correlation between xCT expression and sensitivity to glucose deprivation. As expected, sensitive cell lines (U2OS and U251MG) expressed high levels of xCT compared to intermediate sensitive and resistant cell lines (SW480, MCF7, A375, H1299 and HEK293T), which indicates that xCT could be a biomarker to predict sensitivity to glucose deprivation. However, normal fibroblasts (WI-38 and IMR-90) that are resistant to glucose deprivation, expressed high levels of xCT (Fig. 7A). The expression levels of glucose transporter 1 (GLUT1) and glycolytic enzymes such as hexokinase 1 and 2 and glucose-6-dehydrogenase did not correlate to sensitivity to glucose deprivation. Next, to determine whether xCT expression contributes to NADPH depletion under glucose deprivation, we measured intracellular NADPH levels using multiple cell lines. Sensitive cell lines (U2OS and U251MG) showed rapid NADPH depletion under glucose deprivation, whereas intermediate sensitive and resistant cell lines (SW480, A375, H1299, WI-38 and IMR-90) managed to maintain NADPH levels under glucose deprivation although partial NADPH decrease was observed in some cell lines (Fig. 7B). This indicates a strong correlation between xCT expression, NADPH depletion and sensitivity to glucose deprivation in cancer cells. Only normal fibroblasts (WI-38 and IMR-90) did not show any correlation between xCT levels and NADPH maintenance. A possible explanation is that these normal fibroblasts may have an alternative NADPH source besides glucose-derived-PPP. Further investigation will be needed to verify this.

**Figure 7.**
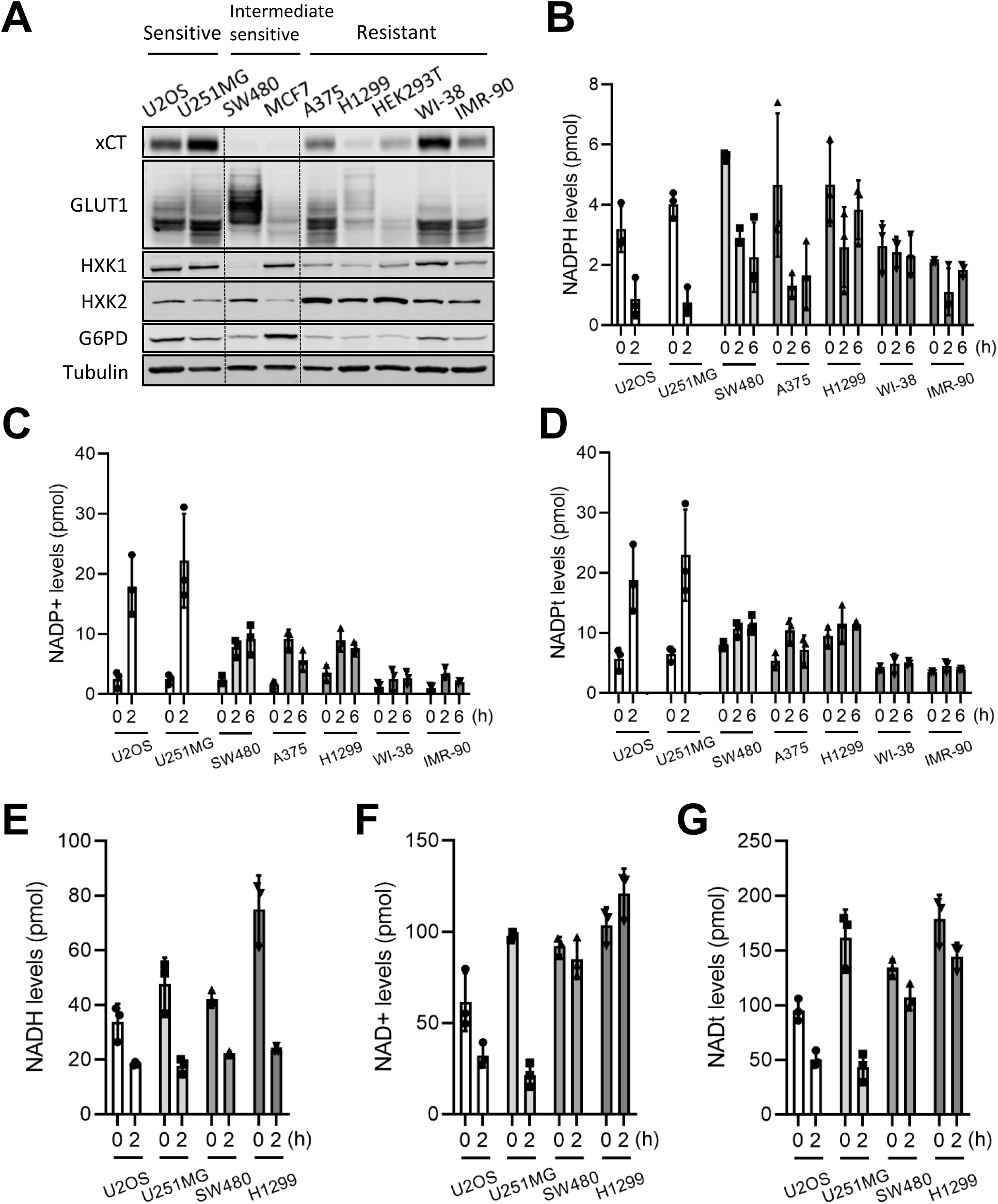
Rapid NADPH depletion under glucose deprivation in cancer cells with high cystine/glutamate antiporter xCT expression. (A) Western blotting analysis was performed using the indicated cell lines. Multiple cell lines were cultured without glucose for the indicated time and intracellular NADPH (B), NADP+ (C) and NADPt (the sum of NADPH and NADP+) (D), NADH (E), NAD+ (F) and NADt (the sum of NADH and NAD+) (G) levels were measured. The mean percentage ± SD of three independent experiments is shown. Images of Western blotting analysis are representative of three independent experiments.

Interestingly, sensitive cell lines (U2OS and U251MG) showed an abnormal increase (4-5 folds) of NADPt (the sum of NADPH and NADP+) which was due to abnormal accumulation of NADP+, whereas intermediate sensitive and resistant cells lines maintained relatively similar levels of NADPt upon glucose deprivation (Fig 7C and D). Given that NADP+ is synthesized from NAD+^50^, we measured intracellular NADH and NAD+ levels upon glucose deprivation in sensitive cell lines (U2OS and U251MG) and resistant cell lines (SW480 and H1299). While both sensitive and resistant cell lines showed decreased NADH levels upon glucose deprivation (Fig. 7E), only sensitive cell lines (U2OS and U251MG) showed decreased NAD+ and NADt (the sum of NADH and NAD+) levels (Fig. 7F and G). NAD+ depletion and NADP+ accumulation after glucose deprivation in sensitive cell lines can be due to increased activity of NAD kinase, NADK that catalyzes the conversion from NAD+ to NADP+^50^. This possibility remains to be investigated.

Taken together, these findings indicate that rapid NADPH depletion upon glucose deprivation is a distinguished feature of sensitive cancer cell lines expressing high levels of xCT. Also, intracellular NADPH deletion is a crucial metabolic determinant of glucose deprivation-induced cell death.

### xCT expression determines sensitivity to glucose deprivation

These data prompted us to investigate whether xCT expression levels determine the glucose dependency of cancer cells. We knocked down xCT with two different siRNA in U2OS cells and monitored sensitivity to glucose deprivation. Knocking down xCT prevented glucose deprivation-induced cell death (Fig. 8A and B) and mitochondrial ROS accumulation (Fig. 8C) and partially rescued NADPH depletion (Fig. 8D). Besides, knocking down xCT restored AMPK activation, reduced mTORC1 activity (Fig. 8E) and prevented glucose withdrawal-induced AMPK oxidation under glucose deprivation (Fig. 8F).

**Figure 8.**
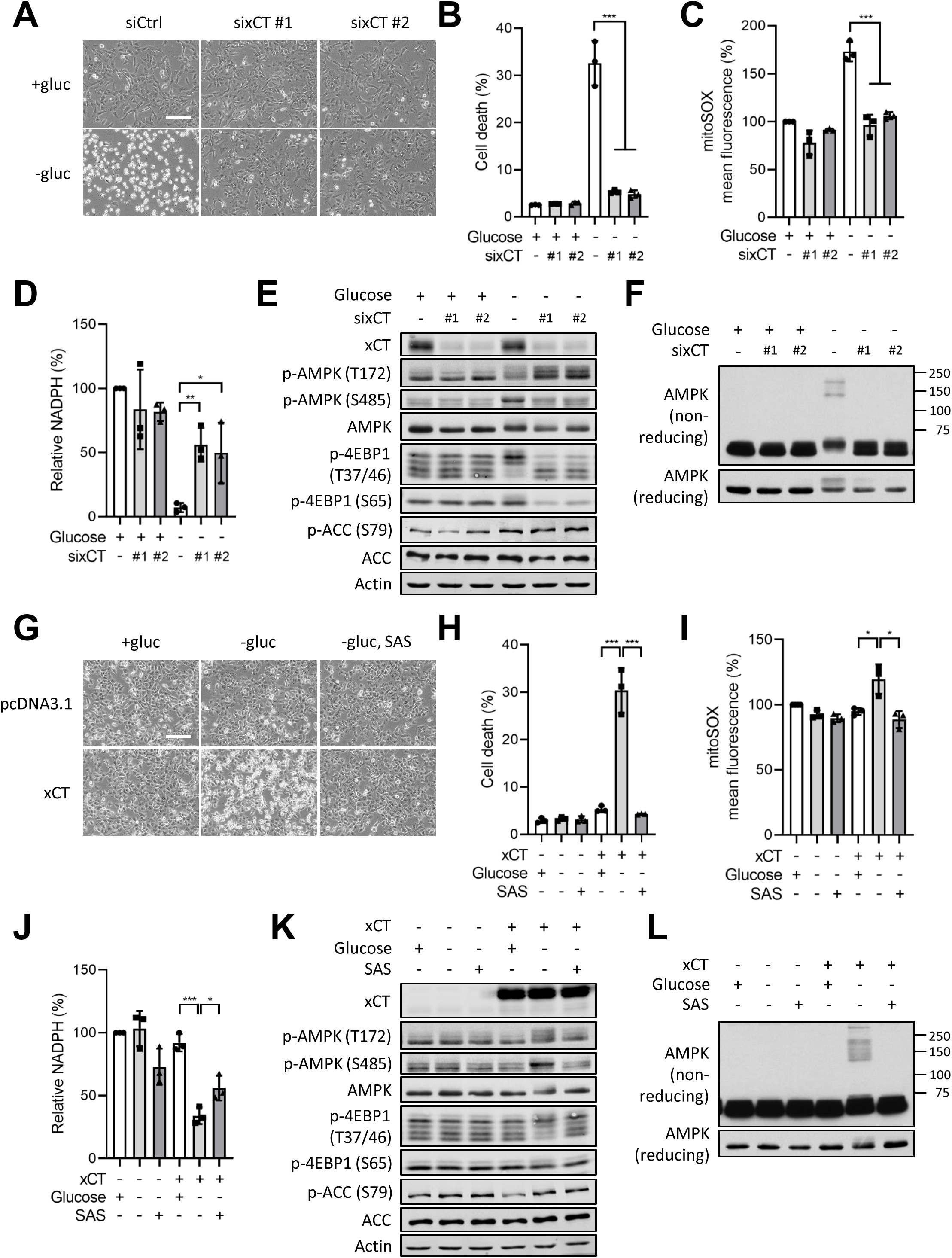
Expressions of cystine/glutamate antiporter xCT determine glucose dependency. siRNA-transfected U2OS cells were cultured in media with or without 1 mM glucose for 3 hours and representative images were taken using phase-contrast microscopy (A); Propidium iodide (PI) exclusion assay was performed at 9 hours (B); mitochondrial ROS was measured at 3 hours (C); intracellular NADPH levels were measured at 2 hours (D); Western blotting analysis was performed at 3 hours (E); non-reducing or reducing Western blotting analysis was performed at 3 hours (F). Empty vector or xCT-overexpressed H1299 cells were cultured in media with or without 1 mM glucose and together with or without 250 μM sulfasalazine (SAS) for 3 hours and representative images were taken using phase-contrast microscopy (G); PI exclusion assay was performed at 9 hours (H); mitochondrial ROS was measured at 3 hours (I); intracellular NADPH levels were measured at 2 hours (J); Western blotting analysis was performed at 3 hours (K); non-reducing or reducing Western blotting analysis was performed at 3 hours (L). Scale bars, 500 μm. The mean percentage ± SD of three independent experiments is shown. Images of Western blotting analysis are representative of three independent experiments. Unpaired two-tailed Student’s t-test was performed for statistical analysis, *P<0.05, **P<0.01, ***P<0.001.

Next, we transiently overexpressed FLAG-xCT in H1299 cells, a resistant cell line to investigate whether high expression of xCT confers high glucose dependency. Overexpression of xCT in H1299 cells potentiated glucose deprivation-induced cell death (Fig. 8G and H), mitochondrial ROS accumulation (Fig. 8I), and NADPH depletion (Fig. 8J), which were prevented by SAS treatment, xCT inhibitor (Fig. 8G, H, I and J). Furthermore, glucose deprivation in xCT overexpressing H1299 cells induced AMPK dysregulation, as shown by increased Ser485 inhibitory phosphorylation of AMPK and mobility shift of total AMPK and increased mTORC1 activity, as represented by phosphorylation of 4EBP1 (Fig. 8K). In non-reducing SDS-PAGE, glucose deprivation-induced AMPK oxidation was only observed in xCT overexpressing H1299 cells, not in empty vector-transfected H1299 cells (Fig. 8L). These phenotypes were prevented by xCT inhibition with SAS treatment (Fig. 8K and L).

Taken together, our results confirmed that xCT expression levels determine sensitivity to glucose deprivation and could be a biomarker to predict therapeutic response to glycolysis-targeting cancer therapy.

### Cystine uptake through xCT is a determinant of GLUT1 inhibition-induced cancer cell death

Our results revealed that cystine uptake renders cancer cells more susceptible to glucose deprivation by accelerating NADPH consumption. These data prompted us to investigate whether glucose dependency could be therapeutically targeted. Because it is physiologically impossible to completely deplete glucose for cancer cells in vivo, we employed STF-31, an inhibitor of the GLUT1^51^. We cultured a sensitive cell line (U2OS), a resistant cell line (H1299) in 2 mM glucose which is within physiological concentration range in the tumor microenvironment^22^ and treated each with STF-31 for 24 hours and evaluated cell death (Fig. 9A and B). We found that STF-31 treatment induced moderate levels of cell death in U2OS cells. But, when treated in combination with additional cystine, GLUT1 inhibitor-induced cell death was significantly sensitized. Furthermore, inhibition of xCT with SAS prevented STF-31-induced cell death completely, which indicates that cell death induced by glucose uptake inhibition is dependent on cystine uptake through xCT. By contrast, STF-31 treatment with or without cystine did not markedly induce cell death in H1299. These data suggest that the glucose dependency of cancer cells is targetable with GLUT1 inhibitor and cystine and that high uptake of cystine through xCT could be a key predictor of sensitivity to GLUT1 targeting therapy.

**Figure 9.**
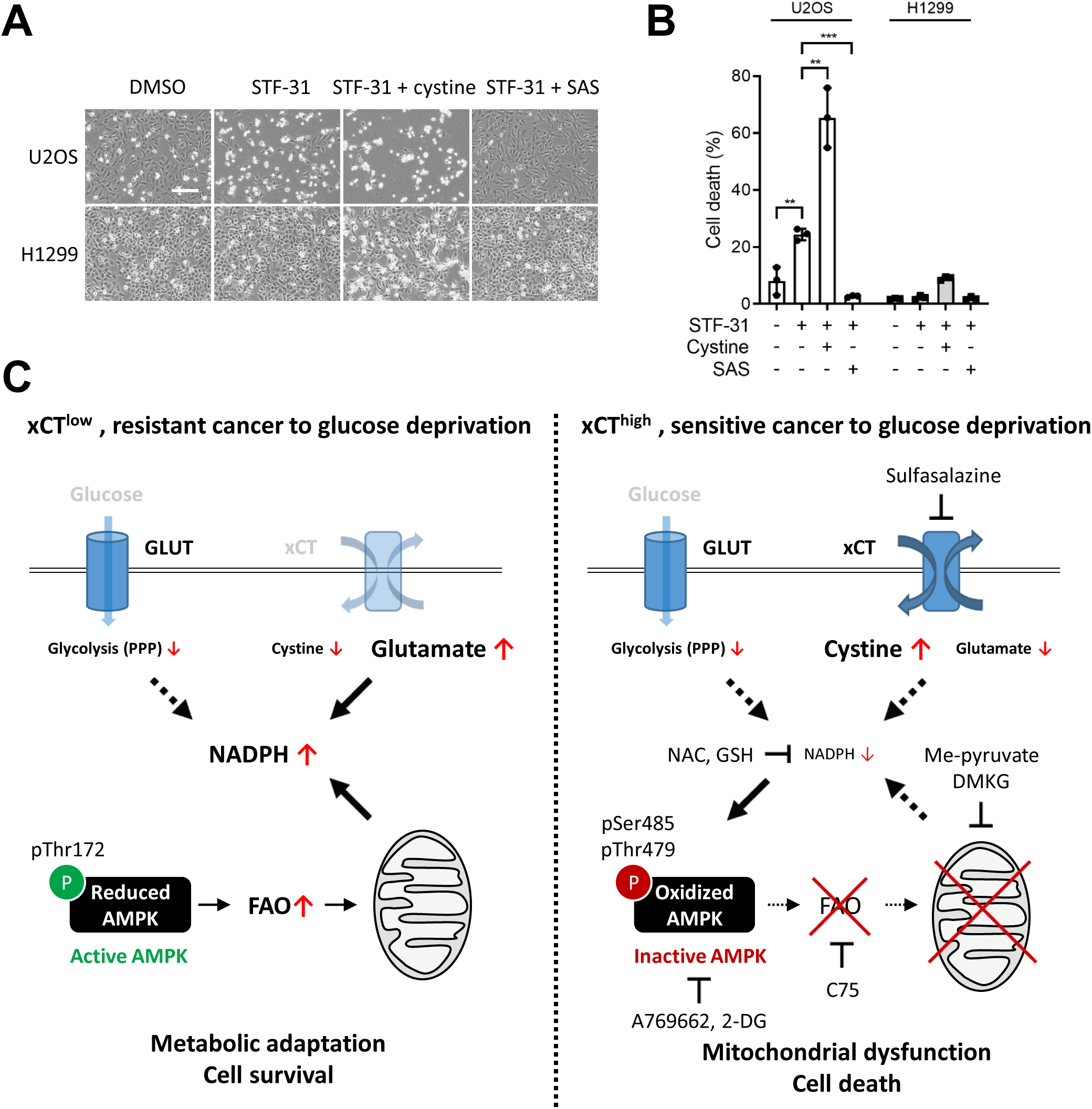
Cystine uptake through cystine/glutamate antiporter xCT is required for GLUT1 inhibitor-induced cell death. U2OS or H1299 cells were cultured in media with 2 mM glucose for 24 hours and 10 μM STF-31, 200 μM cystine or 250 μM sulfasalazine (SAS) was treated simultaneously. Representative images were taken using phase-contrast microscopy (A) and PI-exclusion assay was performed (B). (C) Schematic model of the different regulation of AMPK signaling upon glucose deprivation in low or high xCT-expressing cancer cells. GLUT, glucose transporter; xCT, cystine/glutamate antiporter xCT; PPP, pentose phosphate pathway; FAO, fatty acid oxidation; NAC, N-acetyl-cysteine; GSH, reduced glutathione; DMKG, dimethyl α-ketoglutarate. Scale bars, 500 μm. The mean percentage ± SD of three independent experiments is shown. Unpaired two-tailed Student’s t-test was performed for statistical analysis, **P<0.01, ***P<0.001.

## Discussion

Reprogrammed glucose metabolism has received attention due to possibility of the selective targeting cancer cells. However, the outcome of clinical trials has not been successful due to our limited understanding of glucose deprivation-induced cell death. For example, ATP depletion induced by glucose deprivation was considered the underlying mechanism of cell death and therefore, the glycolysis inhibitor 2-DG was considered for therapy to deplete cellular ATP. However, 2-DG treatment has been shown to prevent glucose deprivation-induced cell death in glucose-sensitive cancer cells despite its ATP-depleting effect^3, 4, 34, 52^, arguing that further study was needed to understand the underlying mechanism of cell death.

Here, we extend our understanding of glucose deprivation-induced cell death with multiple pieces of evidence: a) glucose deprivation sensitive cancer cell lines failed to activate AMPK due to redox-dependent inhibitory oxidation and phosphorylation, leading to failure to adapt to metabolic stress; b) cystine/glutamate antiporter xCT expression is a key biomarker which determines and predicts sensitivity to glucose deprivation; c) NADPH depletion upon glucose deprivation is the key metabolic determinant of glucose deprivation-induced cell death (Fig. 9C).

AMPK is a major metabolic checkpoint that senses energy or nutrient depletion and regulates metabolic homeostasis to maintain cell viability. Therefore, tight regulation of the AMPK pathway is essential for cancer cells to overcome a limited nutrient environment. In the present study, the cell lines most sensitive to glucose deprivation (U2OS and U251MG cells) displayed failure of AMPK activation under metabolic stress, resulting in metabolic catastrophe. LKB1-null cancers are unable to respond to energetic stress due to this lack of an activating kinase for AMPK^4^. Interestingly, although the cell lines we used in the present study have intact LKB1-AMPK axis, these sensitive cells failed to maintain AMPK-mediated adaptive response upon glucose deprivation.

In a glucose-limited environment, cancer cells have to switch their main energy source from glucose to other sources such as glutamine or fatty acid or regulate energy expenditure such as growth inhibition and autophagy induction to overcome metabolic stress. AMPK activation enables cancer cells to overcome metabolic stress by different means. For example, AMPK-mediated resistance to anoikis and nutrient starvation is through inhibition of mTORC1-dependent global mRNA translation^16, 20^. In the present study, although highly sustained mTORC1 activity was observed under glucose deprivation in sensitive cancer cell lines due to defective AMPK activation, mTORC1 inhibitor, rapamycin did not rescue glucose deprivation-induced cell death, indicating high ATP expenditure or mRNA translation due to the high mTORC1 activity is not the main cause of glucose deprivation-induced cell death at least in the cell lines we studied.

A previous study reported that the key difference between sensitive and resistant cells in a glucose-limited environment is the mitochondrial response to metabolic stress and proper mitochondrial OXPHOS becomes essential for cancer cell growth and survival^23^. Particularly, AMPK-dependent metabolic switching to OXPHOS is crucial for cancer cell survival under glucose deprivation. For example, the AMPK-dependent acceleration of FAO renders cancer cells resistant to glucose deprivation by providing Acetyl-CoA, a substrate for the TCA cycle. This process activates mitochondrial OXPHOS, providing an alternative source of ATP and NADPH^4, 53^. Also, AMPK facilitates the TCA cycle by direct phosphorylation of pyruvate dehydrogenase, which improves cancer cells’ adaptation to the metastatic environment for breast cancers^19^. AMPK regulates T cell metabolic adaptation by glutamine-dependent mitochondrial metabolism in a glucose-limited environment^54^. In contrast, LKB1-deficient cancers fail to overcome glucose deprivation or anoikis due to the inability to metabolically switch to FAO^4^, suggesting a major role of AMPK in the metabolic flexibility of cancer cells. In this study, cancer cells with high sensitivity to glucose deprivation likely failed to survive due to the inability to metabolically switch to FAO based on the following: AMPK activators A769662 and 2-DG rescued glucose deprivation-induced cell death, mitochondrial ROS accumulation and NADPH depletion; and fatty acid synthase inhibitor C75 that also facilitates FAO, mimicked the effect of AMPK activators on glucose deprivation-induced cell death. Given that decreased ATP under glucose deprivation was not the main initiator of cell death, AMPK activation is likely to improve cancer cell viability by regulating redox homeostasis via FAO-mediated NADPH production.

The intracellular oxidizing condition can induce covalent disulfide bond formation within and between proteins via cysteine residues, which alters biological functions, subcellular localization and protein-protein interactions^37, 38^. Recent evidence suggests that only specific cysteine residues are susceptible to oxidative stress-induced modification, indicating protein oxidation is a tightly regulated post-translational modification^55^. Here, we found that glucose deprivation in the sensitive cancer cells dampened AMPK-mediated adaptive response by a redox-dependent mechanism. AMPK has been reported to be susceptible to oxidative stress^40, 56^. However, the functional role of AMPK oxidation is debatable depending on the context^40, 57^.

Direct AMPK oxidation by ROS occurs at Cys130 and Cys174 residues, forming intermolecular disulfide bonds and AMPK aggregation, which inhibits AMPK function^40^. In our study, glucose deprivation-induced AMPK dysregulation was rescued by anti-oxidant treatments with NAC or GSH, indicating that oxidative stress caused dysregulation of AMPK. In addition, glucose deprivation induced AMPK mobility shift and aggregation in non-reducing SDS-PAGE, indicating that AMPK undergoes oxidation via the formation of intra- and intermolecular disulfide bonds which interrupt proper AMPK activation. Furthermore, the extent of AMPK oxidation was following redox collapse by NADPH depletion, rather than elevation of cytosolic ROS. This was supported by two lines of evidence: 1) elevation of cytosolic ROS alone by BSO treatment was not sufficient to induce AMPK oxidation. But, diamide treatment which depletes the NADPH pool by oxidizing thiol-residues, was sufficient to induce AMPK oxidation similarly to glucose deprivation in sensitive cell lines; 2) SAS, A769662, C75, me-pyruvate and DMKG treatments which ameliorated mitochondrial ROS accumulation and NADPH depletion but not cytosolic ROS prevented glucose deprivation-induced AMPK oxidation and cell death. A possible explanation is that temporal ROS elevation can activate cellular stress response such as through AMPK^57^. However, severe oxidative stress induced by NADPH depletion can collapse thiol anti-oxidant systems such as TXN and GSH which require NADPH for their recycling. Therefore, this condition can disrupt protein dithiol/disulfide balance, thereby inducing AMPK disulfide bond formation^40^. Why the increase of mitochondrial ROS contributes to AMPK dysregulation and cell death under glucose deprivation is remained to be elucidated. Investigating subcellular localization of NADPH and AMPK under glucose deprivation would be needed to solve this issue.

AMPK has multiple phosphorylation sites which include Thr172 at active loops and Ser485 and Thr479 in the ST loop. Previous studies suggest that phosphorylation in the ST loop negatively regulates the catalytic activity of AMPK by increasing the accessibility of phosphatases to Thr172 phosphorylation^45^ or by restraining LKB1 or CaMKKII-mediated Thr172 phosphorylation^27^. Multiple kinases were reported to phosphorylate serine/threonine residues in the ST loop of AMPK. Akt^27, 42^, PKC^43^ and S6K^44^ can phosphorylate the Ser485 residue. GSK3 phosphorylates Thr479 and its multiple sequential phosphorylation sites, but it requires a preceding Ser485 phosphorylation^28, 45^. Here, we found that glucose deprivation induced an abnormal electrophoretic mobility shift of AMPK due to abnormal phosphorylation including Ser485 in the ST loop. By kinase screening, we found that PKC and GSK3 were responsible for the phosphorylation of Ser485 and other residues in the ST loop, respectively. Although we could not detect Thr479 phosphorylation of AMPK due to the unavailability of the antibody, we suspect that GSK3 might be responsible for Thr479 and subsequence phosphorylation of AMPK based on a previous study^28^.

Interestingly, PKC or GSK3 inhibitor treatment did not rescue glucose deprivation-induced cell death or increase phosphorylation of AMPK substrates such as mTORC1 and ACC although they restored Thr172 phosphorylation of AMPK. In addition, AMPK was still oxidized under this condition. This might be because the inhibitors were not sufficiently able to overcome the rapid NADPH depletion caused by glucose deprivation and thus AMPK activation was insufficient, suggesting that inhibitory oxidation of AMPK is hierarchically dominant than the phosphorylation status of AMPK in preventing activation. Oxidation-induced AMPK aggregation may result in hindering substrate accessibility onto AMPK and so uncoupling it from its substrates.

Unlike in 0 mM glucose, PKC and GSK3 inhibitors rescued cell death by restoring Thr172 phosphorylation and preventing oxidation of AMPK in low glucose (1mM), suggesting that activities of kinases such as PKC and GSK3 hinder proper AMPK activation by inducing abnormal phosphorylation in the ST loop. The difference in results observed in 1 mM and 0 mM glucose might be because NADPH depletion occurs much slowly in 1 mM glucose than 0 mM. Therefore, AMPK activation induced by PKC or GSK3 inhibitors was able to overcome redox collapse and cell death in the 1 mM glucose condition. Why or how PKC and GSK3-mediated phosphorylation of AMPK occurred under glucose deprivation remains puzzling. One possibility is that glucose deprivation-induced oxidative stress may directly or indirectly activate PKC or GSK3 through the regulation of kinases^58^ or phosphatases^3, 6^ respectively.

Recently, multiple studies have demonstrated that xCT activity is required for glucose deprivation-induced cell death. The suggested underlying mechanisms have been various such as mitochondrial dysfunction by glutamate export^59, 60^, NADPH depletion and ROS accumulation by cystine uptake^61, 62^, and disulfide stress^63^. Consistent with these reports, we found that cystine was required for glucose deprivation-induced NADPH depletion, mitochondrial dysfunction and cell death. In particular, our data point to NADPH consumption as a key metabolic determinant of sensitivity to glucose deprivation and the expression levels of xCT determines NADPH consumption and cell death under glucose deprivation. Imported cystine through xCT is quickly reduced to cystine, which consumes NADPH in the process. At the same time, glutamate export through xCT can lead to a reduction of α-ketoglutarate, a metabolite of glutamate entering the TCA cycle where NADPH is produced. Both cystine uptake and glutamate export seem to contribute to the rapid depletion of NADPH under glucose deprivation because replenishing the TCA cycle with DMKG, a cell-permeable α-ketoglutarate was only able to partially rescue the NADPH depletion and cell death. Therefore, high cystine uptake also contributes to NADPH consumption under glucose withdrawal by the cystine-cysteine conversion process.

In addition, sensitive cancer cell lines (U2OS and U251MG) expressed high levels of xCT, compared to resistant cancer cell lines and showed dampened AMPK adaptive pathway to metabolic stress. This AMPK dysregulation was also due to high xCT activity which collapses the redox system under glucose deprivation. High expression of xCT can contribute to glucose deprivation-induced mitochondrial dysfunction in two different ways. First, glutamate export through xCT can cause a shortage of TCA cycle substrate. Second, NADPH depletion-mediated AMPK dysregulation leads to failure of metabolic switching from glycolysis to FAO and shortage of TCA cycle substrate. These data provide insight into a previously underappreciated novel crosstalk between metabolic regulation and signaling transduction which is essential for metabolic adaptation.

Intracellular NAD+ is essential for glycolysis and mitochondrial OXPHOS^50^. Targeting NAD+ metabolism can be therapeutically beneficial to eradicate cancers by inducing metabolic catastrophe^64, 65^. Here, we found that glucose deprivation-induced NAD+ depletion in xCT-high expressing cancer cells. Although the role of NAD+ depletion in glucose deprivation-induced cell death remained to be studied, it could hamper mitochondrial OXPHOS under glucose withdrawal or recovery process responding to glucose re-addition. Therefore, glycolysis-targeting strategy in xCT-high expressing cancers could be effective not only because it causes catastrophic oxidative stress induced by NADPH depletion, but also it may cause metabolic catastrophe due to NAD+ depletion. This possibility remains to be investigated.

In conclusion, our data revealed paradoxical AMPK inactivation in sensitive cancer cell lines expressing high levels of xCT and the underlying molecular mechanism. Given that xCT-high expressing cancer cells are resistant to conventional radio or chemotherapy due to a high generation rate of GSH^47, 49^, a glucose uptake inhibiting strategy together with elevating tissue cystine concentration could be therapeutically more effective for these types of cancers, and xCT expression could be considered as a biomarker for predicting sensitivity to the therapy.

## Methods and materials

### Cell cultures and reagents

U2OS, U251MG, SW480, MCF7, A375, H1299 and HEK293T cells were purchased from American Type Culture Collection (ATCC). WI-38 and IMR-90 cells were purchased from Coriell Institute. All cell lines were cultured in high-glucose Dulbecco’s modified Eagle’s medium (DMEM) (25 mM glucose, glutamine, Gibco, Life Technologies) supplemented with 10% FBS (HyClone, GE Healthcare Life Science), penicillin (100 units/ml) and streptomycin (100 μM/ml; Gibco, Life Technologies) in 5% CO_2_-humidified atmosphere at 37°C. For glucose deprivation, cells were washed with phosphate-buffered saline (PBS) three times and cultured in glucose-free DMEM with 10% dialyzed FBS. DMEM-AA was generated following the recipe of DMEM and glutamine was added.

Glucose, 2-DG, rapamycin, PI, EAA, NEAA, glutamate, oligomycin, FCCP, rotenone, antimycin A, methyl-pyruvate, dimethyl α-ketoglutarate (DMKG), BIO, N-acetyl-cysteine (NAC), reduced glutathione (GSH) were purchased from Sigma-Aldrich. Sulfasalazine (SAS), buthionine sulfoximine (BSO) and A769662 were purchased from Cayman Chemical.

Necrosulfonamide and STF-31 were purchased from Merck Millipore. Deferoxamine, MK-2206 dihydrochloride and C75 were purchased from MedChemExpress. Z-VAD-FMK and diamide are purchased from Santa Cruz Biotechnology. Necrostatin-1 was purchased from Abcam. PKC-412, GF 109203X and Ro 31-8220 mesylate were from Screen-Well^®^ Kinase inhibitor Library Version 2.2 (Enzo Life Sciences). Arginine, cystine, histidine, isoleucine, leucine, lysine, methionine, phenylalanine, threonine, tryptophan, tyrosine and valine were kindly provided by Dr. Jean-Paul Kovalik (Duke-NUS Medical School, Singapore).

### Dialyzed serum

The commercial dialyzed serum (HyClone, GE Healthcare Life Science) was further dialyzed using Slide-A-Lyzer G2 dialysis cassette (MWCO 10,000) (Thermo Fisher Scientific). Dialysis was performed in a cold PBS buffer containing 1 mM phenylmethylsulfonyl fluoride (PMSF) for 24 hours in a cold room.

### RNA interference & plasmids

siRNAs targeting xCT (SLC7A11) were obtained from Qiagen. Control siRNA (ON-TARGET plus nontargeting pool) was obtained from Dharmacon. The targeted siRNA sequences were: xCT #1, AACCACCTGTTTCACTAATAA; xCT #2, TGGGTGGAACTCCTCATAATA. Full-length xCT was amplified from U251MG cells and cloned into vector pcDNA3.1^(+)^ (Thermo Fisher Scientific) with C-terminal 3xFLAG tagging. The cloned construct was confirmed by direct DNA sequencing. Cells were transiently transfected with siRNA or plasmid using ScreenfectA according to the manufacturer’s instructions (FUJIFILM Wako Pure Chemical Corporation). All siRNAs were used at 20 nM for transfection.

### Western blotting analysis

Cells were lysed with 2% SDS lysis buffer (50 mM Tris-HCl, pH 6.8, 10% glycerol and 2% SDS). Equal amounts of proteins (20-30 µg) were subjected to SDS-PAGE. After all proteins were transferred to a nitrocellulose membrane, immunoblots were probed with the indicated antibodies. Detection was done by incubation of HRP-conjugated anti-mouse or anti-rabbit IgG (Jackson ImmunoResearch) secondary antibody followed by the reaction for chemiluminescence (SuperSignal,Thermo Fisher Scientific). Infrared fluorescence-conjugated anti-mouse or anti-rabbit IgG secondary antibody (Dy-light, Jackson ImmunoResearch) was used for infrared fluorescence detection (LI-COR Odyssey).

For AMPK redox analysis, cells were washed with ice-cold PBS once and lysed with 0.1% NP-40 lysis buffer (50 mM Tris-HCl pH 7.5, 150 mM NaCl, 1 mM PMSF, 1x protease inhibitor cocktail without EDTA from WAKO, 1x phosphatase inhibitor cocktail from Roche) containing 20 mM N-Ethylmaleimide to prevent further cysteine oxidation during protein extraction. Equal amounts of proteins (20 µg) were subjected to SDS-PAGE with loading buffer with or without DTT and β-ME. The dotted line in Western blotting images does not indicate separated images. The following antibodies were used; anti-AMPK (Cell Signaling Technology, #2793), anti-phospho-AMPK Thr172 (Cell Signaling, #2535), anti-phospho-AMPK Ser485 (Cell Signaling, #4185), anti-ACC (Cell signaling, #3676), anti-phospho-ACC Ser79 (Cell Signaling, #11818), anti-phospho-Akt Thr308 (Cell Signaling, #9275), anti-G6PD (Cell Signaling, #8866), anti-LC3B (Cell Signaling, #2775), anti-LKB1 (Cell Signaling, #3047), anti-phospho-S6K Thr389 (Cell Signaling, #9234), anti-xCT/ SLC7A11 (Cell Signaling, #12691), anti-phospho-4EBP1 Ser65 (Cell Signaling, #9451), anti-phospho-4EBP1 Thr37/46 (Cell Signaling, #9459), anti-GLUT1 (abcam, #ab115730), anti-HXK1 (Santa Cruz, #sc-6517), anti-HXK2 (Santa Cruz, #sc-6521), anti-actin (Merck Millipore, clone C4, cat # MAB1501) anti-tubulin (Abcam, clone no. DM1A+DM1B).

### PI exclusion assay

Cells were stained with PI to determine the percentage of cell death. Media containing floating cells were collected, combined with trypsinized cells, and centrifuged. The cell pellet was washed once with PBS. After centrifugation, cells were re-suspended and stained with PI (10 μg/mL) for 10 mins at room temperature. Data were collected with MACSQuant analyzer (Miltenyi Biotec). Quantification and analysis of the data were done with Flowjo software.

### **λ** phosphatase assay

Cells were lysed with 0.5% NP-40 lysis buffer (0.5% NP-40, 50 mM Hepes, pH 7.5, 100 mM NaCl, 1x protease inhibitor cocktail without EDTA from WAKO), and endogenous phosphatase activity was heat-inactivated at 75°C for 5 mins. A total of 400 units of λ phosphatase (New England Biolabs) was added to each lysate and incubated at 30°C for 1 hour. Enzyme activity was heat-inactivated at 95°C for 5 mins before Western blotting analysis.

### ATP measurement assay

Cells (2×10^4^/ well) were seeded in 96 well plates. Intracellular ATP levels were measured by CellTiter-Glo luminescent cell viability assay (Promega) according to the manufacturer’s guideline.

### Cytosolic and mitochondrial ROS measurement assay

Cells were deprived of glucose for 3 hours. During the last 30 mins of glucose deprivation, 2.5 μM DCFDA or 2.5 μM mitoSOX (Thermo Fisher Scientific) was added to media for staining for 30 mins at 37°C. Media containing floating cells were collected, combined with trypsinized cells, and centrifuged. The cell pellet was washed once with PBS. Data were collected with MACSQuant analyzer (Miltenyi Biotec). Quantification and analysis of the data were done with Flowjo software.

### NADPH and NADH measurement assay

Intracellular NADPH/NADP+ and NADH/NAD+ were measured using the NADP/NADPH Quantification Kit (ab65349, Abcam) and NAD/NADH Quantification Kit (ab65348, Abcam) according to the manufacturer’s instructions. Briefly, 2.5×10^5^ cells were lysed with 350 μL of extraction buffer at indicated time points after glucose deprivation. For the reaction, 50 μL of final sample was used. Signal Intensities for NADPH or NADH were examined by OD measurements at 450 nm using Infinite M200 plate reader (TECAN).

### GSH/GSSG measurement assay

Intracellular GSH/GSSG was measured using GSH/GSSG-Glo^TM^ luminescent assay (Promega) according to the manufacturer’s instructions. Briefly, 2 x10^4^ cells in 96 well plates were lysed at the indicated time point in the indicated condition.

### Mitochondrial oxygen consumption rate (OCR) measurement

OCR was measured using a Seahorse Bioscience XF96 Extracellular Flux Analyzer (Seahorse Bioscience). 2 x 10^4^ cells were plated into Seahorse tissue culture 96 well plates. Cells were cultured in DMEM with or without 1 mM glucose for 2 hours. And then cells were cultured in Seahorse assay media containing 2 mM glutamine with or without 1 mM glucose and incubated in a CO_2_-free incubator for an hour before measurement. XF Cell Mito Stress Test Kit was used to analyze mitochondrial metabolic parameters by measuring OCR. Oligomycin (1 μM) was injected to determine the oligomycin-independent lack of the OCR. The mitochondrial uncoupler FCCP (1 μM) was injected to determine the maximum respiratory capacity. Rotenone (1 μM) and antimycin A (1 μM) were injected to block complex I and complex III of the electron transport chain.

### Statistical analysis

Data are presented as mean ± SD of at least three independent experiments. Unpaired two-tailed Student’s t-test was performed for statistical analysis. All statistical analyses were conducted using GraphPad Prism 9.0.2 software.

## Acknowledgments

We thank Drs. David M Virshup, Anne-Claude Gingras, Egon Ogris, Brijesh K Singh and Jennifer Alagu for the discussion of the data and the manuscript. This work was supported by Duke-NUS Signature Programme Block Grant and the Singapore Ministry of Health’s National Medical Research Council grants (NMRC/OFIRG/15nov049/2016) to K.I.

## Author contribution

Y.L. and C.C.O. performed experiments. Y.L., Y.I. and K.I. designed the experiments and analyzed the results. Y.L., Y.I. and K.I. wrote the manuscript.

## Conflict of Interest

No potential conflicts of interest were disclosed by all the authors.

---

**Supplementary figure S1.**
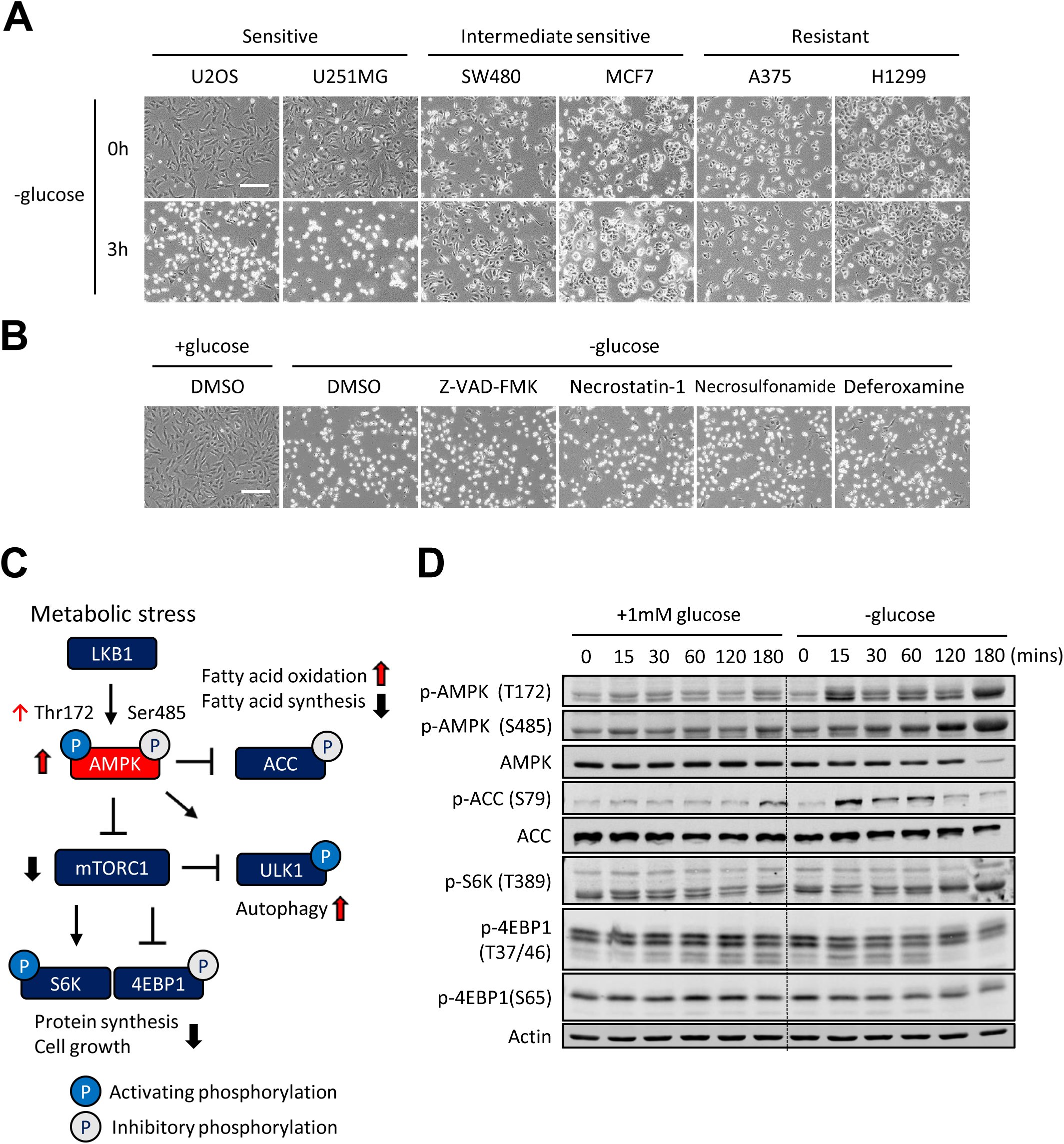
(A) The indicated cell lines were cultured in media without glucose for 3 hours and representative images were taken using phase-contrast microscopy. (B) U2OS cells were cultured in media with or without 1 mM glucose for 3 hours and representative images were taken using phase-contrast microscopy. The indicated cell death inhibitor was treated simultaneously. (C) Schematic model of general changes of AMPK-mTORC1 signaling under metabolic stress and functional consequences. (D) Time-course analysis of Western blotting was performed using U251MG cells cultured with or without 1 mM glucose for the indicated time. Scale bars, 500 μm.

**Supplementary figure S2.**
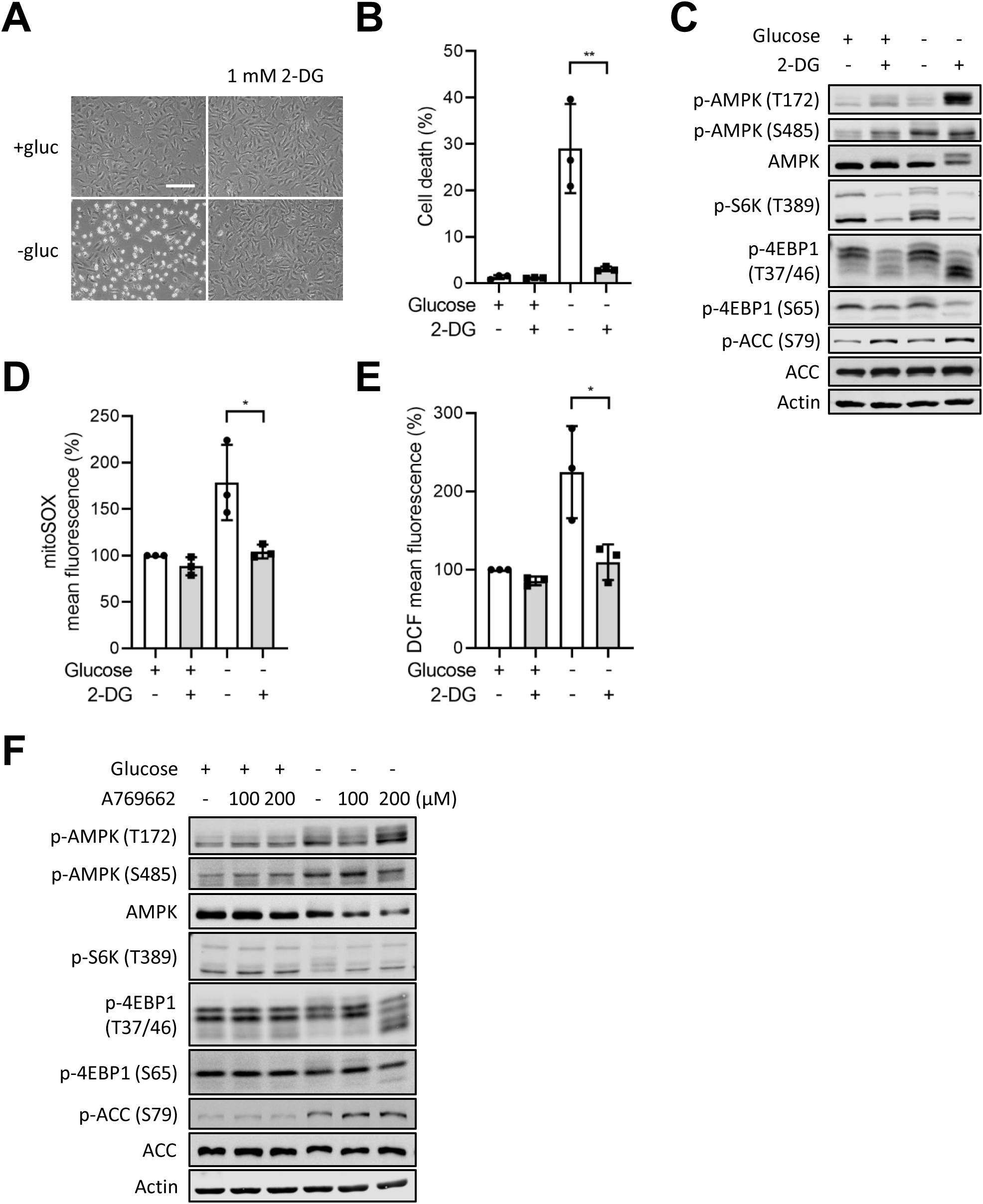
U2OS cells in media with or without 1 mM glucose were treated with 1 mM of 2-deoxyglucose (2-DG) for 3 hours and representative images were taken using phase-contrast microscopy (A); Propidium iodide (PI) exclusion assay was performed after 9 hours (B); Western blotting analysis was performed at 3 hours (C); mitochondrial (D) and cytosolic (E) ROS was measured at 3 hours. (F) U251MG cells in media with or without 1 mM glucose were treated with the indicated doses of A769662 for 3 hours and harvested for Western blotting analysis. Scale bars, 500 μm. The mean percentage ± SD of three independent experiments is shown. Images of Western blotting analysis are representative of three independent experiments. Unpaired two-tailed Student’s t-test was performed for statistical analysis, *P<0.05, **P<0.01.

**Supplementary figure S3.**
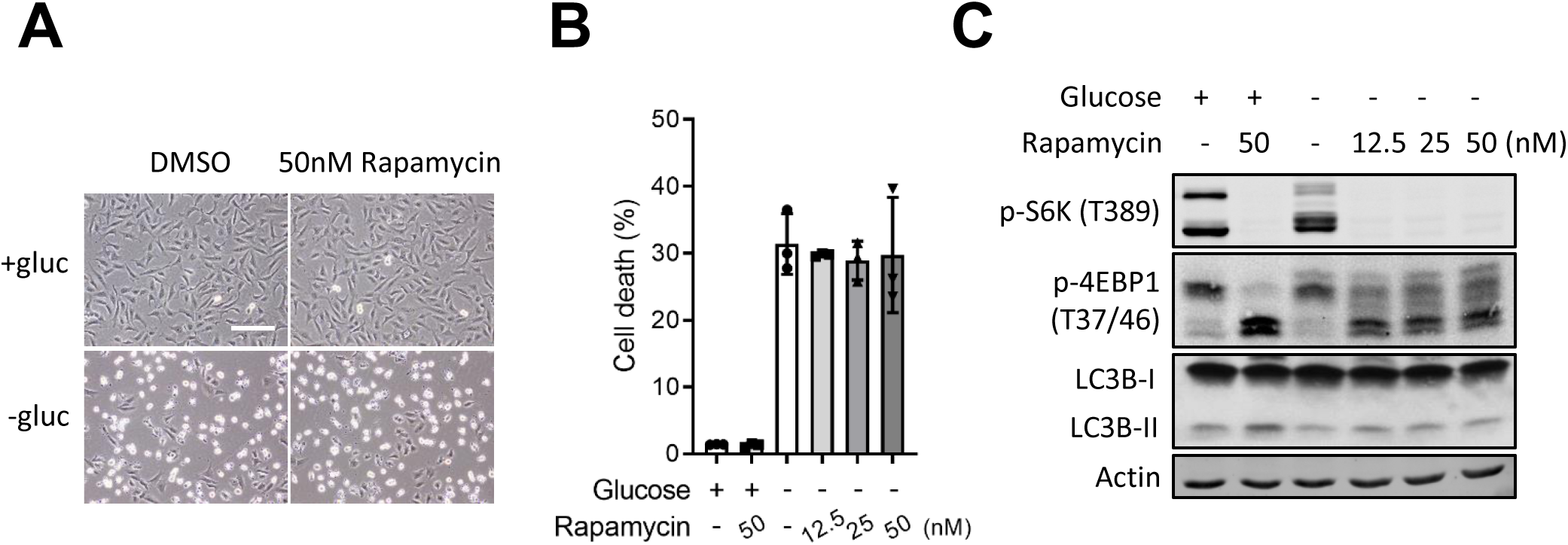
U2OS cells were pre-treated with the indicated dose of rapamycin for 1 hour and then cultured in media with or without 1 mM glucose for 3 hours. Rapamycin was treated simultaneously. Representative images were taken using phase-contrast microscopy (A); PI exclusion assay was performed after 9 hours (B); Western blotting analysis was performed at 3 hours (C). Scale bars, 500 μm. The mean percentage ± SD of three independent experiments is shown. Images of Western blotting analysis are representative of three independent experiments.

**Supplementary figure S4.**
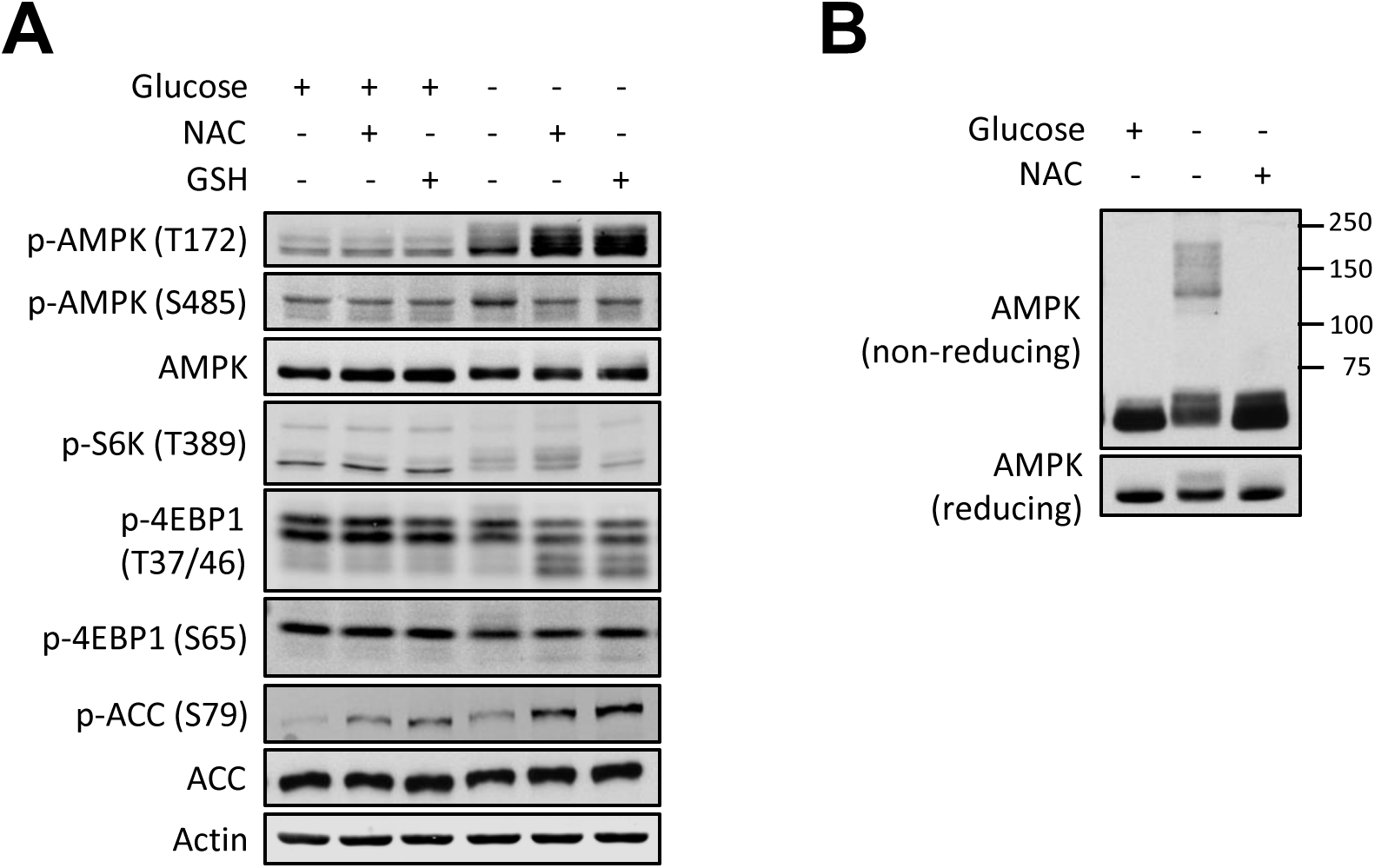
(A) U251MG cells in media with or without 1 mM glucose were treated with the 5 mM N-acetyl-cysteine (NAC) or reduced glutathione (GSH) for 3 hours and harvested for Western blotting analysis. (B) U251MG cells in media with or without 1 mM glucose were treated with the 5 mM NAC for 3 hours and harvested for non-reducing or reducing Western blotting analysis. Images of Western blotting analysis are representative of three independent experiments.

**Supplementary figure S5.**
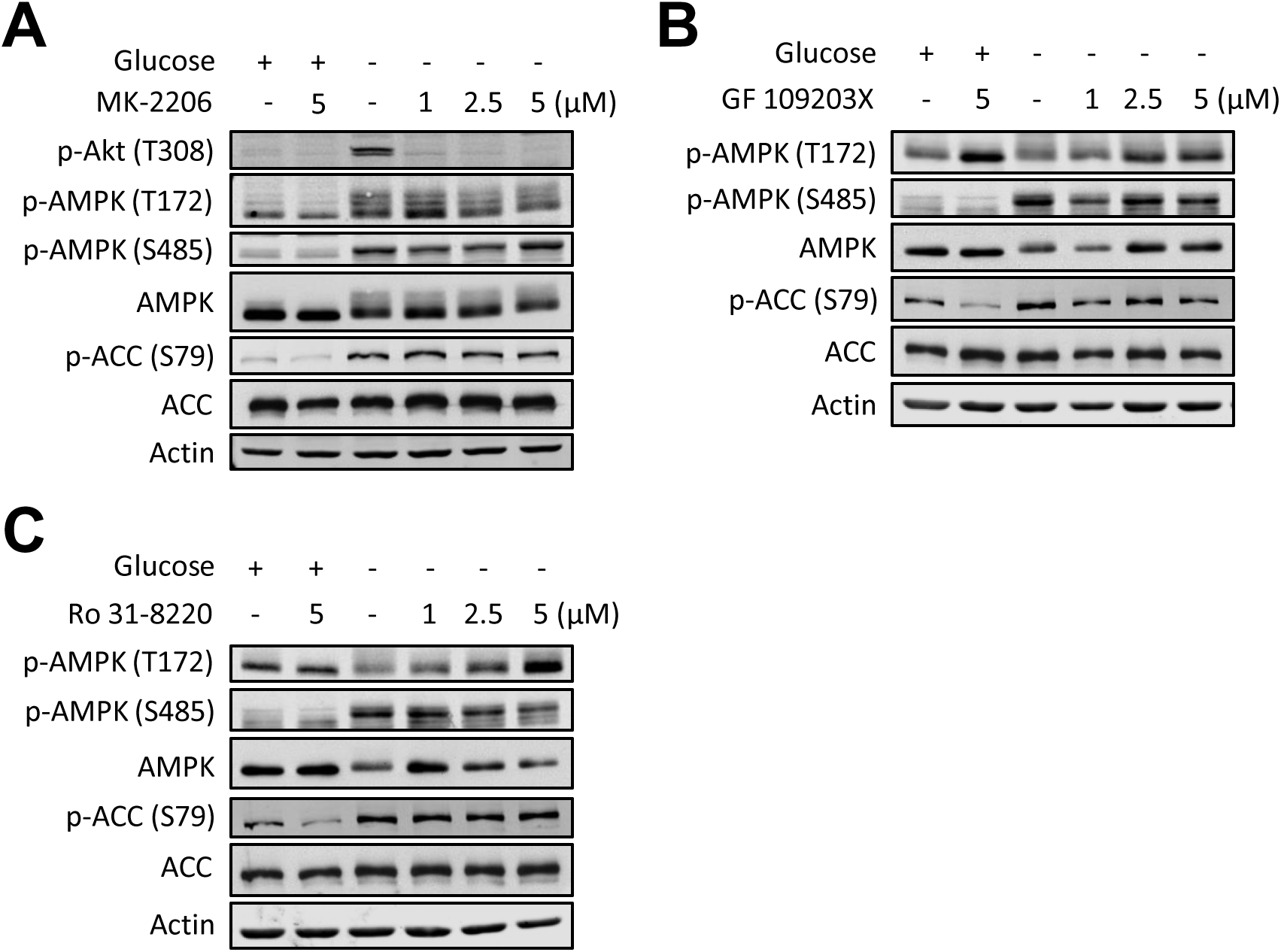
U2OS cells in media with or without 1 mM glucose were treated with the indicated doses of MK-2206 (A), GF 109203X (B) or Ro 31-8220 (C) for 3 hours and harvested for Western blotting analysis. Images of Western blotting analysis are representative of three independent experiments.

**Supplementary figure S6.**
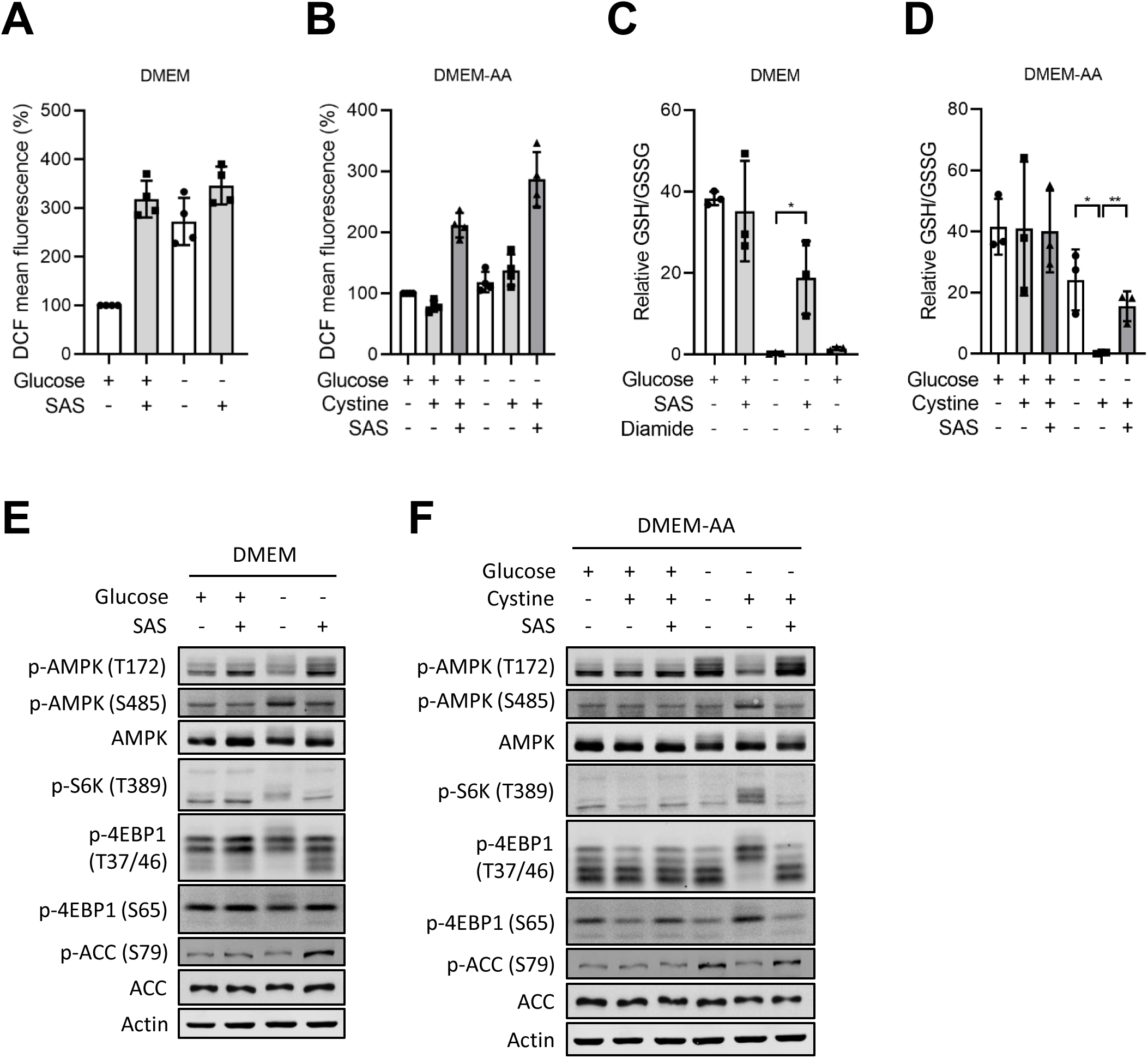
U2OS cells were cultured in the DMEM (A) or amino acid-free DMEM (DMEM-AA) (B) with or without 1 mM glucose for 3 hours. 250 μM sulfasalazine (SAS), 200 μM cystine or 250 μM diamide was treated as indicated simultaneously. Cytosolic ROS was measured at 3 hours (A and B); intracellular reduced glutathione (GSH)/ oxidized glutathione (GSSG) ratio was measured at 2 hours (C and D); Western blotting analysis was performed (E and F). Images of Western blotting analysis are representative of three independent experiments. The mean percentage ± SD of three or more than three independent experiments is shown. Unpaired two-tailed Student’s t-test was performed for statistical analysis, *P<0.05, **P<0.01.

## Notes

### Competing Interest Statement

The authors have declared no competing interest.

